# Genomic structural variation in clownfish adaptive radiation

**DOI:** 10.64898/2026.09.25.754321

**Authors:** Anna Marcionetti, Ema Inacio Martins, Diego A. Hartasánchez, Fabio Cortesi, Lucy M. Fitzgerald, Bruno Frederich, Theo Gaboriau, Alberto García-Jiménez, Vincent Laudet, Sarah Schmid, Nicolas Salamin

## Abstract

Clownfishes are a complex of 28 species that rapidly diversified after acquiring mutualism with host sea anemones around 15 million years ago. The genomic mechanisms behind this radiation are only beginning to emerge, and structural variants (SVs) have received little attention. Previous phylogenomic studies suggested two large inversions on chromosome 18, but their distribution across the clownfish phylogeny and their potential effects have not been formally explored. SVs can disrupt gene function and regulation or alter gene dosage, and they have been shown to play a central role in adaptive evolution and diversification across many taxa.

Here, we characterized SVs across the clownfish radiation using long-read PacBio sequencing, generating high-quality genome assemblies for sixteen species spanning the main lineages of the genus. We identified and catalogued nearly 130,000 SVs spanning close to 570 Mb. Short insertions and deletions were the most numerous, but large inversions accounted for most of the affected sequence. No SV was consistently associated with host specialization. We nevertheless confirmed a single inversion of approximately 17 Mb on chromosome 18, clarifying earlier reports of two separate rearrangements at this locus, and identified an additional inversion on chromosome 9 whose distribution suggests a history of hybridization.

This study provides the first genome-wide characterization of structural variation in clownfishes, revealing extensive SVs despite the group’s rapid and recent radiation. It also delivers new, high-quality genomic resources that will support future research into this iconic group of coral reef fishes and their adaptation to sea anemones.

## Introduction

Clownfishes (genus *Amphiprion*) are an iconic group of coral reef fishes distributed across the Indo-Pacific Ocean, known for their obligate mutualism with sea anemones. This mutualism originated 10 to 15 million years ago and is thought to have acted as a key innovation that triggered the adaptive radiation of the group (Litsios et al., 2012). Following the acquisition of the ability to live among the toxic tentacles of sea anemones, clownfishes rapidly diversified into 28 recognized species showing convergence in colouration and morphology, driven by specialization to different host sea anemone species (Litsios et al., 2012; Gaboriau et al., 2025). Host associations range from generalists inhabiting up to ten sea anemone species to strict specialists restricted to a single host genus, namely *Radianthus*, *Entacmaea*, or *Stichodactyla* (Gaboriau et al., 2025). Differences in swimming-related traits have also been documented across species and may have further contributed to the diversification of the group (Mercader et al., 2025).

Numerous studies have investigated the molecular mechanisms underlying clownfish diversification. How clownfishes tolerate the toxic tentacles of sea anemones has been examined through biochemical and genomic approaches (Nedosyko et al., 2014; Marcionetti et al., 2019; Roux et al., 2019; Heim et al., 2025; Roux et al., 2025; Suresh et al., 2026), and the genomic basis of host specialization and convergent phenotypic evolution has also been explored (Marcionetti and Salamin, 2023; Heim et al., 2023; Gaboriau et al., 2025; Fitzgerald et al., 2026b). Despite these advances, the precise mechanisms driving clownfish diversification remain unclear. Genes showing convergent positive selection across clownfish species associated with specialization to *Radianthus* or *Entacmaea* hosts have been identified (Gaboriau et al., 2025), but causal links between specific variants and phenotypic divergence have yet to be determined. Hybridization has also been proposed as a contributor to clownfish diversification (Litsios and Salamin, 2014; Schmid et al., 2022; Marcionetti and Salamin, 2023; Schmid et al., 2025), and phylogenomic analyses have revealed a striking pattern of topological inconsistency on chromosome 18, hypothesized to reflect large chromosomal inversions suppressing recombination across that region (Schmid et al., 2022; Marcionetti and Salamin, 2023; Marcionetti et al., 2024). While these inversions have not been formally confirmed, these findings suggest that genomic structural variation may have played an important role in clownfish diversification. Yet, such variation has never been systematically characterized in the group.

Genomic structural variants (SVs) are large genomic rearrangements (*>* 50 bp) that include insertions, deletions, duplications, inversions, translocations, and chromosomal fusions. These variants represent a major source of genomic and phenotypic variation across taxa. Unlike single nucleotide variants, SVs can simultaneously affect large genomic regions, disrupting or duplicating entire genes, reshaping regulatory landscapes, and altering gene dosage, often with substantial functional consequences (Mérot et al., 2020; Zhang et al., 2021). The importance of SVs in local adaptation and species diversification has become increasingly evident across a wide range of organisms (Wellenreuther et al., 2019; Mérot et al., 2020). For example, copy number variation (CNV) is responsible for variation in plumage colouration and thermal adaptation in the common murre (*Uria aalge*, Tigano et al., 2018), and Robertsonian fusions involving sex chromosomes contribute to reproductive isolation in the killifishes (genus *Lucania*, Berdan et al., 2021b). Large inversions, often spanning several Mbp, are also frequently associated with adaptive phenotypes (Mérot et al., 2020; Wellenreuther et al., 2025). By suppressing recombination between inverted and standard haplotypes, inversions can maintain sets of locally adaptive alleles in strong linkage disequilibrium, thereby forming “supergenes”. These regions facilitate the coordinated inheritance of complex adaptive traits and allow selection to act more efficiently, even in the face of ongoing gene flow (Mérot et al., 2020; Wellenreuther et al., 2025). This process has been documented in diverse systems, with well-known examples including the mimicry supergene of *Heliconius* butterflies (Joron et al., 2011) and the supergene underlying alternative reproductive strategies in the ruff (*Philomachus pugnax*; Küpper et al., 2016). Inversions have also been documented to facilitate speciation and reproductive isolation in adaptive radiations (Zhang et al., 2021; Combrink et al., 2025), with notable examples in sticklebacks (Jones et al., 2012) and cichlids (Blumer et al., 2025).

Given the growing evidence for the role of SVs in species diversification, we aimed to investigate the extent and evolutionary significance of structural variation in clownfishes. The advent of long-read sequencing technologies, which avoid problems associated with short-reads, such as the erroneous collapse of duplicates (Hartasánchez et al., 2018), have made it possible to characterize SVs in evolutionary-diverse clades such as *Amphiprion*. Using HiFi long-read sequencing, we generated high-quality reference genomes for 16 clownfish species, bringing the total number of clownfish reference genomes to 18 out of the 28 known species. Using these resources, we set out to characterize structural variation across the clownfish clade to explore the association between SVs and specialization to particular sea anemone hosts.

## Materials and Methods

### Species selection, DNA extraction, library preparation and sequencing

Fin clips for 16 clownfish species (*Amphiprion akallopisos*, *A. akindynos*, *A. allardi*, *A. biaculeatus*, *A. chrysopterus*, *A. ephippium*, *A. frenatus*, *A. latifasciatus*, *A. latezonatus*, *A. mccullochi*, *A. omanensis*, *A. percula*, *A. perideraion*, *A. polymnus*, *A. sandaracinos*, *A. sebae*) were obtained through fieldwork expeditions (research permit reported in García-Jiménez et al., 2025), collaborators at the Marine Station at the Academia Sinica (Taiwan), and through aquarium shops (Supplementary Table S1).

Species were selected to cover the full clownfish divergence and to encompass multiple independent transitions in sea anemone host specialization (specialization to *Entacmaea* hosts: *A. omanensis*, *A. mccullochi*, *A. frenatus*, *A. ephippium*, *A. biaculeatus*; specialization to *Radianthus* hosts: *A. akallopisos*, *A. perideraion*, *A. ocellaris* and *A. percula*; specialization to *Stichodactyla* hosts: *A. polymnus* and *A. sandaracinos*; Figure 1A; (Gaboriau et al., 2025).

**Figure 1:**
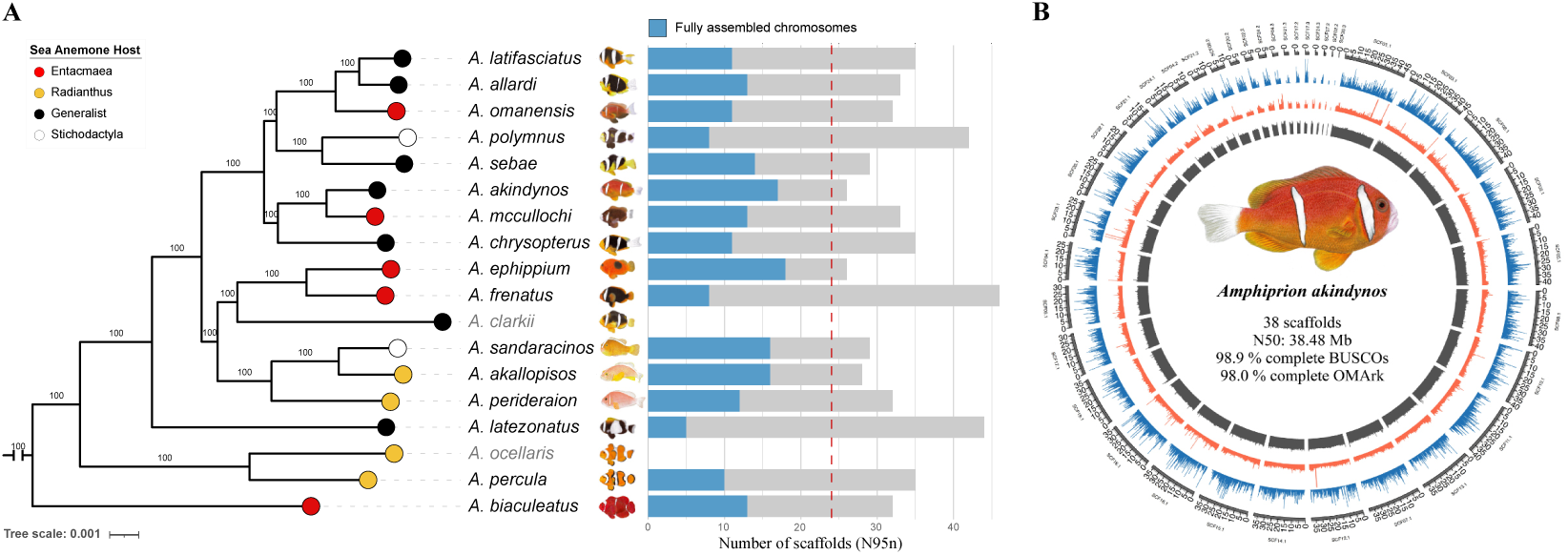
**A)** Phylogenetic relationships among the clownfish species assembled in this study (labels in black) and publicly available reference genomes (*Amphiprion clarkii* GCA 027123335.1 and *A. ocellaris* GCA 022539595.1, in grey). Associations with host sea anemones, as reported in Gaboriau et al., 2025, are indicated by coloured circles. For the species assembled in this study, the number of resulting scaffolds is reported as N95n (i.e. the number of scaffolds covering 95% of the assembly length). The blue portion of each bar corresponds to fully assembled chromosomes, and the vertical red dotted line indicates the expected number of chromosomes in the genus (24 chromosomes). The phylogenetic tree was inferred using IQ-TREE2 (v2.2.2.7; Minh et al., 2020) on 14,733 single-copy orthologs from clownfishes and the damselfishes *Acanthochromis polyacanthus* (GCA 021347895.1) and *Dascyllus trimaculatus* (GCA 024666655.1), which were used for rooting and are not shown in the final tree. Branch support values are shown above branches. The tree and barplot were visualised using iTOL (v7.5.1; Letunic and Bork, 2024). **B)** Genome assembly and annotation overview for *Amphiprion akindynos*, shown as a representative example of the genome assemblies generated in this study. The outer circle represents the assembled scaffolds. Tracks display gene density (blue), repeat density (red), and GC content (grey). The number of scaffolds, N50 length, genome assembly completeness (BUSCO, Simão et al., 2015) and annotation completeness (OMArk, Nevers et al., 2025) are reported at the center of the plot. The circular plot was generated using Circos (Krzywinski et al., 2009).

For each species, high-molecular-weight genomic DNA was extracted from approximately 100 mg of fin tissue following standard DNA extraction protocols. The total amount of DNA was measured with the Qubit dsDNA HS Assay Kit, and integrity was verified using the Fragment Analyzer Automated CE System. DNA fragments larger than 10 kb were selected using the BluePippin System (Sage Science, USA), and HiFi libraries were generated using the SMRTbell Template Prep Kit 2.0 (PacBio, USA) at the Genomic Technology Facility (GTF) in Lausanne. Seven species were sequenced using Circular Consensus Sequencing (CCS) on a PacBio Sequel II system (one lane per species), while the remaining species were sequenced on a PacBio Revio system (three species per lane across three lanes).

### Genome assembly

HiFi read quality was assessed using FastQC (v0.12.1) and summarized with MultiQC (v1.25.1; Ewels et al., 2016). Genome characteristics were examined by counting k-mers with Jellyfish (v2.2.7; Marçais and Kingsford, 2011) and GenomeScope (v1.0; Vurture et al., 2017). To determine the most suitable genome assembler for our data, we evaluated four software packages: the Improved Phased Assembler (IPA) from PacBio (v1.8.0; https://github.com/PacificBiosciences/pbipa), Flye (v2.9.1; Kolmogorov et al., 2019), Hi-Canu (v2.2; Nurk et al., 2020) and Hifiasm (v0.19.5; Cheng et al., 2022). Assemblies were generated for four species (*A. akallopisos*, *A. ephippium*, *A. perideraion*, *A. polymnus*). The quality and completeness of the assemblies were assessed using `assembly-stats` (v1.0.1; https://github.com/sanger-pathogens/assembly-stats) and BUSCO (v5.4.7; actinopterygii odb10 database; Simão et al., 2015). Assembly accuracy was further evaluated by aligning HiFi reads back to the assemblies with minimap2 (v2.12; Li, 2018) and computing alignment statistics using samtools (v1.19.2; Li et al., 2009).

Hifiasm and HiCanu produced the most complete assemblies (Supplementary Table S2); however, with HiCanu, the resulting assembly sizes were approximately twice the expected genome sizes, likely indicating haplotig duplication. Duplicated haplotypes were removed using purge dups (v1.2.5; Guan et al., 2020), and the assemblies were sub-sequently re-evaluated for completeness and accuracy. Although this procedure improved HiCanu assembly metrics, Hifiasm consistently outperformed the other softwares and was thus chosen as the best assembler for our data (Supplementary Table S2).

The genomes of the 12 additional species were assembled using Hifiasm (v0.19.5; Cheng et al., 2022) and potential haplotypic duplications were removed with purge dups (v1.2.5; Guan et al., 2020). For all species, further filtering was performed: mitochondrial sequences and duplicated scaffolds (defined as sequences with at least 80% of their length matching other scaffolds at more than 80% identity) were identified using MUMmer4 (v4.0.0; programs `nucmer`, `delta-filter` and `show-coords`; Marçais et al., 2018) and removed with custom scripts (https://github.com/amarcion/ClownfishSV/).

The number of chromosomes (24) appears conserved in clownfish, as shown by the chromosome-level reference genomes of *A. clarkii* (GCA 027123335.1, Moore et al., 2023), *A. percula* (GCA 003047355.2; (Lehmann et al., 2019)), *A. ocellaris* (GCA 022539595.1; (Ryu et al., 2022)) and *A. biaculeatus* (GCA 053813585.1; (Zhang et al., 2026); see also Whole-genome alignments and structural variation identification section below). As chromosome fusions were not expected, assemblies were screened for potential misassemblies by alignment to the *A. clarkii* reference genome using MUMmer4 (v4.0.0; programs `nucmer`, `delta-filter` and `show-coords`; Marçais et al., 2018). Scaffolds aligning to multiple chromosomes over more than 25% of their length were considered misassembled and split accordingly.

Additional scaffolding steps were performed to further reduce assembly fragmentation. First, scaffolds were clustered by putative chromosomal origin based on *A. clarkii* alignments. When assignments were ambiguous, clusters associated with multiple chromosomes were merged. Scaffolding was then performed independently within each cluster using HiFi reads with SAMBA (MaSuRCA v4.1.0; parameters: -m 2500 -d asm; Zimin and Salzberg, 2022; Zimin et al., 2013) and ntLink (v1.3.9; parameters: -n 2; Coombe et al., 2021). Subsequently, we applied a scaffolding strategy based on P RNA scaffolder (Zhu et al., 2018). For each species, artificial paired-end (PE) libraries (5, 7, 8, 11 and 15 kb inserts) were generated from HiFi reads using custom scripts (https://github.com/amarcion/ClownfishSV/). The PE reads were mapped to the assemblies with BWA (v0.7.17; Li and Durbin, 2009) and the resulting alignments were processed using SAM-tools (v1.19.2; Li et al., 2009). Scaffold links identified by P RNA scaffolder were filtered by coverage (2x-25x) and insert size consistency, merged across libraries, and used for scaffolding with a default gap size of 100 Ns. This pipeline was iterated five times with shifted read extraction positions. Finally, scaffolds shorter than 200 kb were removed.

### Assembly quality

Assembly quality was assessed at each step with Inspector (v1.3.1; Chen et al., 2021), and error correction was performed using the software’s correction module (`inspector-correct.py`). To ensure that the assembly steps did not introduce significant errors, we evaluated the number of structural and small-scale errors and estimated the Quality Value (QV) using the software. Each assembly step reduced scaffold number with minimal impact on QV and error counts (Supplementary Table S3, Supplementary Figure S1). All species exhibited QV values above 43, exceeding those reported in similar studies (e.g.; Krawczyk et al., 2025; Maughan et al., 2024; Parey et al., 2024; Pu et al., 2024). Additional assembly statistics were computed with `assembly-stats` (v1.0.1; https://github.com/sanger-pathogens/assembly-stats) and completeness was estimated using BUSCO (v5.4.7; actinopterygii odb10 database; Simão et al., 2015). Statistics are reported in Supplementary Table S4.

To further characterise final assembly quality and individual heterozygosity, HiFi reads were mapped to the corresponding final assemblies using minimap2 (v2.28; -`x map-hifi`; Li, 2018). SNP variants were called with Clair3 (v2.0.1; parameters `--platform=hifi` and `--model path=hifi sequel2` or **--model path=hifi revio** depending on the sequencing platform; Supplementary Table S1; Zheng et al., 2022), and SVs were called with Sniffles2 (v2.5; Smolka et al., 2024). Because reads originated from the same individuals used for assembly, heterozygous variants reflect within-individual genetic diversity, whereas an excess of homozygous variants or anomalous coverage patterns may indicate residual assembly errors. Sequencing depth, SNPs and SVs statistics were computed using VCFtools (v0.1.16; Danecek et al., 2011) and custom scripts (https://github.com/amarcion/ClownfishSV/). Individual heterozygosity was estimated in non-overlapping 100-kb genomic windows. Windows containing fewer than 50 SNPs were excluded, and the mean and standard deviation of heterozygosity were calculated across the remaining windows.

### Genome annotation

Repetitive elements were modeled using EDTA (v2.1.3; Ou et al., 2019) and Repeat-Modeler2 (v2.0.1; Flynn et al., 2020), including the LTR structural discovery module (-LTRStruc). Resulting repeat libraries were merged and complemented with the DFAM Vertebrata library (release 3.3) and the clownfish-specific repeat library from Marcionetti and Salamin (2023). Assemblies were then soft- and hard-masked using RepeatMasker (v4.0.7; Smit et al., 2015).

Gene annotation was performed on the soft-masked assemblies with BRAKER (v3.0.3; Gabriel et al., 2024) using protein homology evidence from the OrthoDB Eukaryota database (v11; Kuznetsov et al., 2023). Gene models were evaluated by homology searches against the Swiss-Prot database (release 2024 1; The UniProt Consortium et al., 2025) using DIAMOND (v2.0.15; Buchfink et al., 2021). Functional domains were identified with InterProScan (v5.66.98.0; Jones et al., 2014) against the Pfam (v36; Mistry et al., 2021), PANTHER (v18; Mi et al., 2005) and Antifam (v7; Eberhardt et al., 2012) databases. Only gene models supported by Swiss-Prot, Pfam or PANTHER matches and lacking Antifam hits were retained as the final gene set.

Functional annotation was performed using DIAMOND v2.0.15; Buchfink et al., 2021) searches against the NCBI non-redundant protein database (downloaded 2024-02-07). Gene ontology (GO) terms were assigned using the NCBI `gene2accession`, `gene2go` and `gene info` datasets (downloaded 2024-04-02). Annotation quality and completeness were evaluated with BUSCO (v5.4.7; actinopterygii odb10 database; Simão et al., 2015) and OMArk (v0.3.1; Metazoa database; Nevers et al., 2025).

Genome annotation was also performed as reported above for the reference genome of *Dascyllus trimaculatus* (GCA 024666655.1; Roberts et al., 2023), for which a genome assembly was available but no annotation had been provided. Assemblies and annotations were visualized with Circos (v0.69-8; Krzywinski et al., 2009). Gene and repeat densities were calculated with `dsComputeBEDDensity` from deepStats (v0.3.1; Richard, st 6).

### Orthologous gene inference and phylogenetic reconstruction

Hierarchical orthologous groups (HOGs) were inferred using OMA standalone (v2.6.0; Altenhoff et al., 2019). In addition to the proteomes of the newly assembled species and *Dascyllus trimaculatus* (GCA 024666655.1), we included the publicly available proteomes from *A. clarkii* (GCA 027123335.1), *A. percula* (GCA 003047355.2), *A. ocellaris* (GCA 022539595.1), *Acanthochromis polyacanthus* (GCA 021347895.1), *Oreochromis niloticus* (GCA 001858045.3) and *O. aureus* (ASM587006v1). OMA output was processed with PyHAM (v1.2.0; Train et al., 2019) to extract HOGs across taxonomic levels. Custom scripts were used to identify single-copy orthologous genes (1:1 OGs) shared among all clownfish species and at least one damselfish species (*D. trimaculatus* and *A. polyacanthus*; https://github.com/amarcion/ClownfishSV/).

For each 1:1 OG (total of 14,733 OGs), coding sequences were retrieved, aligned using MAFFT (v7.505; Katoh and Standley, 2013), and trimmed with trimAl (v1.4.1; parameters: `-gt 0.75 -st 0.001`; Capella-Gutiérrez et al., 2009). The resulting alignments were concatenated with AMAS (Borowiec, 2016), and species phylogenetic relationships were inferred using IQ-TREE (v2.2.2.7; Minh et al., 2020). A partitioned model was applied, allowing each locus to evolve at its own rate, and branch support was assessed using the SH-like approximate likelihood ratio test and ultrafast bootstrap (parameters: `-m GTR+G+I-alrt 1000 -bb 1000`). The phylogenetic tree was visualized with iTOL (v7.5.1; Letunic and Bork, 2024). The damselfish species (*D. trimaculatus* and *A. polyacanthus*) were used as outgroups for rooting and were subsequently removed from the tree.

### Clownfishes ancestral scaffolds and chromosomal rearrangements

To investigate the overall large-scale structural variation in clownfish genomes, ancestral scaffolds were inferred using AGORA (v3.1; “agora-generic” pipeline; Muffato et al., 2023), with gene families derived from inferred HOGs (total of 40,823 HOGs) as input. Orthologous relationships between extant genomes and the reconstructed clownfish ancestor were compared using AGORA’s src/misc.compareGenomes.py (modes `printOrthologuesList` and `printOrthologousChrom`), considering scaffolds or chromosomes larger than 200 genes. Karyotype plots were visualized with AGORA’s src/misc.compareGenomes.py script in `drawKaryotype` mode and with syntenyPlotteR (v1.0; Quigley et al., 2023).

### Whole-genome alignments and structural variation identification

Whole-genome alignments among available clownfishes reference genomes (*A. biaculeatus* GCA 053813585.1; *A. clarkii* GCA 027123335.1; *A. ocellaris* GCA 022539595.1; *A. percula* GCA 003047355.2) were performed using minimap2 (v2.28; `parameters -x asm5`; Li, 2018). Results were visualized with the script `pafCoordsDotPlotly.R` from dotPlotly (https://github.com/tpoorten/dotPlotly). For *A. percula* and *A. biaculeatus*, new genome assemblies were generated in this study despite the availability of published references. The published *A. percula* assembly, based on older sequencing data, showed extensive rearrangements relative to other chromosome-level genomes (Supplementary Figure S2); therefore, an independent assembly was obtained here and used for downstream analyses. The chromosome-level reference of *A. biaculeatus* was only recently available (Zhang et al., 2026), and thus an independent assembly was generated here. Our *A. biaculeatus* assembly showed no major structural differences relative to the reference (Supplementary Figure S3).

Using the chromosome-level assembly of *A. biaculeatus* (Zhang et al., 2026) as the reference, whole-genome alignments were generated for all species, including publicly available assemblies of *A. ocellaris* and *A. clarkii*. Alignments were again performed with minimap2 and visualized with dotPlotly as described above.

Genomic structural variants (SVs) relative to *A. biaculeatus* were identified from the whole-genome alignments using SyRI (v1.6.3; Goel et al., 2019). As SyRI requires assemblies with matching chromosome numbers, analyses were conducted either on complete chromosome sets (when available) or on the longest scaffolds. SVs outputs were visualized using plotsr (v1.1.1; Goel and Schneeberger, 2022). For each species, the number and cumulative length of each SV class (i.e., inversions, duplications, translocations, insertions, and deletions greater than 50 bp) were calculated relative to *A. biaculeatus* and visualized using ggplot2 (v3.4.4; Wickham, 2016).

SV callsets were indexed with tabix (v1.21; Li, 2011) and merged using vcf-merge from VCFtools (v0.1.13; Danecek et al., 2011) to generate a multi-species VCF. Variants of the same class with start coordinates within 1 kb across species were treated as a single variant.

Principal coordinates analysis (PCoA) based on SV presence-absence was performed on the full dataset and separately for each SV class using the R function cmdscale. Pairwise dissimilarities between species were calculated using the Jaccard distance metric as implemented in the vegan R package (v2.7-3; Oksanen et al., 2026). SV distributions across species, including variant number, cumulative length, and class composition, were summarized and visualized using ggplot2 (v3.4.4; Wickham, 2016).

To assess the robustness of SV inference to reference choice, all analyses were repeated using the chromosome-level *A. clarkii* assembly (Moore et al., 2023) as reference. The impact of the aligner was also evaluated by repeating the full pipeline with MUMmer4 (v4.0.0; Marçais et al., 2018) for both *A. biaculeatus* and *A. clarkii* references, yielding consistent results (data not shown).

### SVs annotation and catalogue

Fo each SV identified, the genomic coordinates were intersected with the *A. biaculeatus* genome annotation (Zhang et al., 2026) using the GenomicRanges (v1.62.1; Lawrence et al., 2013) and rtracklayer (v1.70.1; Lawrence et al., 2009) R packages. SVs overlapping at least one annotated protein-coding sequence (CDS) feature were considered as potentially affecting the protein sequence and function, and the corresponding gene IDs were recorded. As no functional annotation was available for the *A. biaculeatus* reference genome, the proteome was searched against the NCBI non-redundant protein database (downloaded 2024-02-07) using DIAMOND blastp (v2.0.15; Buchfink et al., 2021), retaining the best hit as a proxy for protein function.

The interactive SV catalogue was generated using a custom Python script that parsed the spreadsheet of the identified and annotated SVs using the openpyxl library. Chromosome identifiers were cross-referenced to NCBI accessions. The output is a self-contained HTML file with data embedded as JSON. The interactive table was implemented using DataTables 1.13.8 with the Bootstrap 5.3.2 styling plugin, providing client-side sorting, full-text search, and paginated browsing. Filter controls allow subsetting by SV category (Private/Shared), SV type (DEL, INS, DUP, TRANS, INV), chromosome, species, SV length range, and minimum number of species. Clicking any table row opens a modal panel with full details including affected transcripts, species list, and BLAST protein hits. A one-click CSV export of the filtered view is provided via the DataTables Buttons plugin. The Python generation script and the interactive HTML interface were developed with the assistance of Claude (Anthropic).

### SVs and sea anemone host specialization

To test for an association between structural variation and sea anemone host specialization, species were assigned to one of four host categories following Gaboriau et al. (2025): *Entacmaea* specialists (*n* = 5), *Radianthus* specialists (*n* = 4), *Stichodactyla* specialists (*n* = 2), and generalists (*n* = 6; Fig 1). An SV was considered host-specific when it was present in all members of a given host category and absent from all other species.

To determine whether host-specific SVs were more common than expected by chance, we compared the number of SVs shared within each specialist group to those identified in equally sized control groups composed of species with similar phylogenetic distances but different host associations. For *Stichodactyla* specialists, all three possible control pairs were used: *A. akallopisos*–*A. polymnus*, *A. akallopisos*–*A. sebae*, and *A. sandaracinos*–*A. sebae*. For *Radianthus* specialists, only one suitable control group was available: *A. ocellaris*–*A. percula*–*A. perideraion*–*A. sandaracinos*, in which *A. akallopisos* was replaced by its closest relative *A. sandaracinos*. No shared SVs were identified among *Entacmaea* specialists, excluding equivalent comparisons for this group.

All analyses were conducted in R (v4.5.2; R Core Team, 2025) and figures were generated using ggplot2 (v3.4.4; Wickham, 2016). Repeating the analyses with the SV dataset generated using the *A. clarkii* reference genome produced consistent results (Supplementary FigureS4).

### Long insertions and topological inconsistencies

Among SVs, inversions stand out because they suppress recombination between the standard and inverted haplotypes, allowing linked variants to persist even under ongoing gene flow. Because of this, inversions have been linked to local adaptation and diversification across a range of taxa (Mérot et al., 2020; Wellenreuther et al., 2025). Because recombination is suppressed, inversions can preserve genealogical histories that differ from the rest of the genome, particularly when they have been introduced through hybridization or when ancestral inversion polymorphisms persist through incomplete lineage sorting. As a result, gene trees from inverted regions may reflect shared chromosomal arrangements rather than genome-wide evolutionary relationships, producing local phylogenies that conflict with the species tree. To identify inversions that might be evolutionarily important, we screened the SV catalogue for long inversions and examined whether they generated phylogenetic relationships that reflected chromosomal arrangement rather than genome-wide species tree.

To reconstruct local phylogenetic trees, we first retrieved the SNPs called by SyRI between each species and the *A. biaculeatus* reference genome, keeping only homologous positions falling within syntenic, inverted, or translocated alignment blocks. Using custom scripts (https://github.com/amarcion/ClownfishSV/), we combined the SNP information across species into a single VCF containing all SNP genotypes. For positions falling outside a homologous alignment block in a given species, the SNP genotype was coded as missing data.

We selected inversions longer than 100 kb and present in at least two species. For each inversion, we extracted the corresponding SNP dataset, removed species with more than 60% missing data, and retained only regions with at least nine remaining species, resulting in a final set of 68 inversions. We inferred phylogenetic trees using IQ-TREE (v2.2.2.7; Minh et al., 2020) under a GTR model with ascertainment bias correction, assessing branch support using the SH-like approximate likelihood ratio test and ultrafast bootstrap (-m GTR+ASC -alrt 1000 -bb 1000). For each region, we tested whether species carrying the inversion formed a monophyletic group in the inversion-specific tree despite not being monophyletic in the species tree, using the R package ape (v5.8.1; Paradis and Schliep, 2019). Two inversions, located on chromosomes CM132566.1 and CM132553.1 (Fig. 3), showed this pattern and were selected for further investigation.

For the two inversions of interest (Fig. 3), we assessed whether each inversion was homozygous within each species by examining split-read evidence at the inversion breakpoints. Breakpoint coordinates were identified in each species’ assembly from pairwise alignments with the *A. biaculeatus* reference genome. We retrieved the mapped HiFi reads to each species’ assembly (Section Assembly quality) and, for each breakpoint, we extracted the reads overlapping a 1 kb window centered on the breakpoint coordinate using SAMtools (v1.19.2; Li et al., 2009). We classified reads as continuous or split based on the presence of a supplementary alignment (SA) tag. Reads spanning the breakpoint that mapped continuously supported the reference haplotype, whereas reads mapping as split/supplementary supported an inverted haplotype. The resulting proportion of split reads thus reflects genotype at the breakpoint, with values near 0 or 1 indicating homozygosity and intermediate values (around 0.5) indicating heterozygosity. Read mapping quality at the breakpoints was also visually investigated with `wally` (v0.6.1; Rausch et al., 2023).

We examined repeat content at inversion breakpoints using genome-wide Repeat-Masker annotations (Section Genome annotation). For each breakpoint, we first determined whether it overlapped an annotated repeat. We then tested whether the 5 kb region surrounding the breakpoint was enriched or depleted for repeat content relative to the rest of the chromosome. For this, we calculated the total repeat content (in bp) within a 5 kb window centered on each breakpoint and compared with an empirical background distribution generated by tiling the corresponding chromosome into non-overlapping 5 kb windows and calculating repeat content for each window. Two-tailed empirical p-values were computed as the proportion of background windows with repeat content at least as extreme as that observed at the breakpoint.

We also investigated gene content at the breakpoints and within each inversion using the final protein-coding gene annotations of each species (Section Genome annotation). We examined genes spanning the breakpoints, as well as genes located within 5 kb on either side of each breakpoint, as these may have been affected by the inversion. Functional annotations of protein-coding genes located within each inversion were retrieved and used to perform Gene Ontology (GO) enrichment analyses with the R package TopGO (v2.62.0; Fisher’s exact test, weight01 algorithm, minimum node size = 5; Alexa and Rahnenführer, 2025) for each inversion and species independently. GO terms with P ¡ 0.05 were considered significant. Following the recommendations of the TopGO manual, no correction for multiple testing was applied. We retained GO terms that were significantly enriched in all species. Finally, we manually examined genes within each inversion for functions related to behaviour, morphology, colouration, and ecology, as these traits may have contributed to clownfish diversification.

To evaluate the extent of topological discordance associated with inversions relative to the rest of the genome, we further assessed phylogenetic variation at the chromosome scale. We inferred phylogenetic trees for each chromosome using the same IQ-TREE approach described above, based on SNPs identified across the entire chromosome. We then compared chromosome-level topologies with the whole-genome species tree inferred from HOGs (Fig. 1) by calculating Robinson–Foulds distances in R (v4.5.2; R Core Team, 2025) using the `ape` package (v5.8.1; Paradis and Schliep, 2019). Pairwise topological distances were visualized using classical multidimensional scaling (`cmdscale`), with plots generated using `ggplot2` (v4.0.2; Wickham, 2016).

## Results

### Whole genome sequencing, genome assembly and annotation

Whole-genome sequencing of the 16 clownfish species generated high-quality long-read data, with average read lengths ranging from 9 to 18 kb and sequencing depths between 26X and 54X (Supplementary Table S5). These data allowed the construction of highly contiguous and complete genome assemblies (Fig. 1; Supplementary Table S4; Supplementary Figures S5–S19). Assembly sizes were highly consistent across species, with scaffold numbers ranging from 35 in *A. akallopisos* to 78 in *A. frenatus*. Contiguity was further reflected by N90 counts ranging from 23 (for *A. akindynos* and *A. ephippium*) to 37 (for *A. frenatus* and *A. latezonatus*), corresponding to N90 scaffold lengths between 6 Mb and 20.64 Mb, respectively. The longest scaffold exceeded 40 Mb in every assembly. Comparative analyses against the chromosome-level genome of *A. clarkii* (Moore et al., 2023) suggested that between 5 (*A. latezonatus*) and 18 (*A. ephippium*) out of the expected 24 chromosomes were fully assembled in each species (Fig. 1A).

Assembly contiguity was substantially improved through successive filtering and scaffolding steps (see Genome assembly), while quality values (QVs) and error metrics remained largely unchanged (Supplementary Figure S1; Supplementary Table S3). Final assemblies achieved QV scores above 43 in all species, exceeding values reported in comparable studies (e.g., Krawczyk et al., 2025; Maughan et al., 2024; Parey et al., 2024; Pu et al., 2024). Sequencing depth across assemblies matched expected values (Supplementary Figure S20A), and mapping reads back to the assemblies yielded very few homozygous single nucleotide polymorphisms (SNPs) or homozygous SVs, both of which are potential indicators of assembly errors (Supplementary Figure S20C–D). Together, these metrics support the high accuracy of the final assemblies.

Repeat content ranged from 27.03% in *A. biaculeatus* to 33.77% in *A. chrysopterus* (Supplementary Table S6). DNA transposons represented the most abundant repeat class across all genomes, accounting for an average of 15.4% of assembly length, followed by long interspersed nuclear elements (LINEs; 3.8%) and long terminal repeat retrotransposons (LTRs; 2.5%). These repeat landscapes are broadly consistent with previous studies on clownfish genomes (e.g., Ryu et al., 2022; Moore et al., 2023; Zhang et al., 2026), although the overall abundance of repeats varies between species.

Genome annotation identified between 26,925 and 27,771 protein-coding genes per species (Supplementary Table S7), of which more than 97% were functionally annotated. Annotation completeness was high, with over 92% of BUSCO genes (Simão et al., 2015) recovered and more than 96.6% of conserved HOGs detected using OMArk (Nevers et al., 2025) (Supplementary Table S7; Supplementary Figure S21). Although a small proportion of genes were classified as duplicated, the overall annotation quality was high and comparable to that reported for other clownfish genomes (e.g., Marcionetti et al., 2019; Ryu et al., 2022; Moore et al., 2023; Zhang et al., 2026). The whole-genome phylogeny inferred from 14,733 single-copy orthologous groups (1:1 OGs) recovered the expected relationships among species, confirming the presence of repeated shifts in host usage (Fig. 1).

Because haplotypes were collapsed during assembly, heterozygous variants provide an estimate of within-individual genetic diversity. Genome-wide heterozygosity varied considerably among species, with per-window values ranging from 0.00109 *±*6.43 × 10*^−^*^4^ in *A. biaculeatus* to 0.00497 *±*2.6 × 10*^−^*^3^ in *A. akindynos* (Supplementary Figure S20B). Patterns of heterozygous structural variation were consistent with these estimates, with counts ranging from 13,474 in *A. biaculeatus* to 45,019 in *A. akindynos*. In all species, most heterozygous SVs consisted of short insertions or deletions smaller than 1 kb (Supplementary Figure S22).

### Synteny conservation and overall structural variation across clownfishes

To investigate large-scale chromosomal rearrangements and assess patterns of synteny across clownfish genomes, we first reconstructed ancestral scaffolds and compared them to extant species. We found no evidence of major chromosomal-scale rearrangements such as fusions or fissions (Supplementary Figure S24), and synteny was highly conserved across species. Most ancestral chromosomal blocks were recovered intact despite some assembly fragmentation (Supplementary Figure S24). Nevertheless, local structural variation, including inversions of large syntenic blocks, was present across the phylogeny.

To better quantify this local structural variation, we performed whole-genome alignments of all species against the *A. biaculeatus* chromosome-level reference genome (Zhang et al., 2026) and identified SVs using SyRI (Goel et al., 2019). Consistent with the conserved macro-synteny described above, genome-wide alignments revealed a high degree of sequence conservation across species, with single-nucleotide divergence ranging from 1.51% (*A. akindynos*, and *A. omanensis*) to 1.61% (*A. latezonatus*; Supplementary Table S8). Nevertheless, SVs were detected in all species, with the proportion of the genome affected ranging from 6.3% in *A. perideraion* to 14.3% in *A. percula* (Fig. 2A; Supplementary Table S8). Highly similar results were obtained when *A. clarkii* reference genome (Moore et al., 2023) was used as the reference (Supplementary Figure S23, Supplementary Table S9).

**Figure 2:**
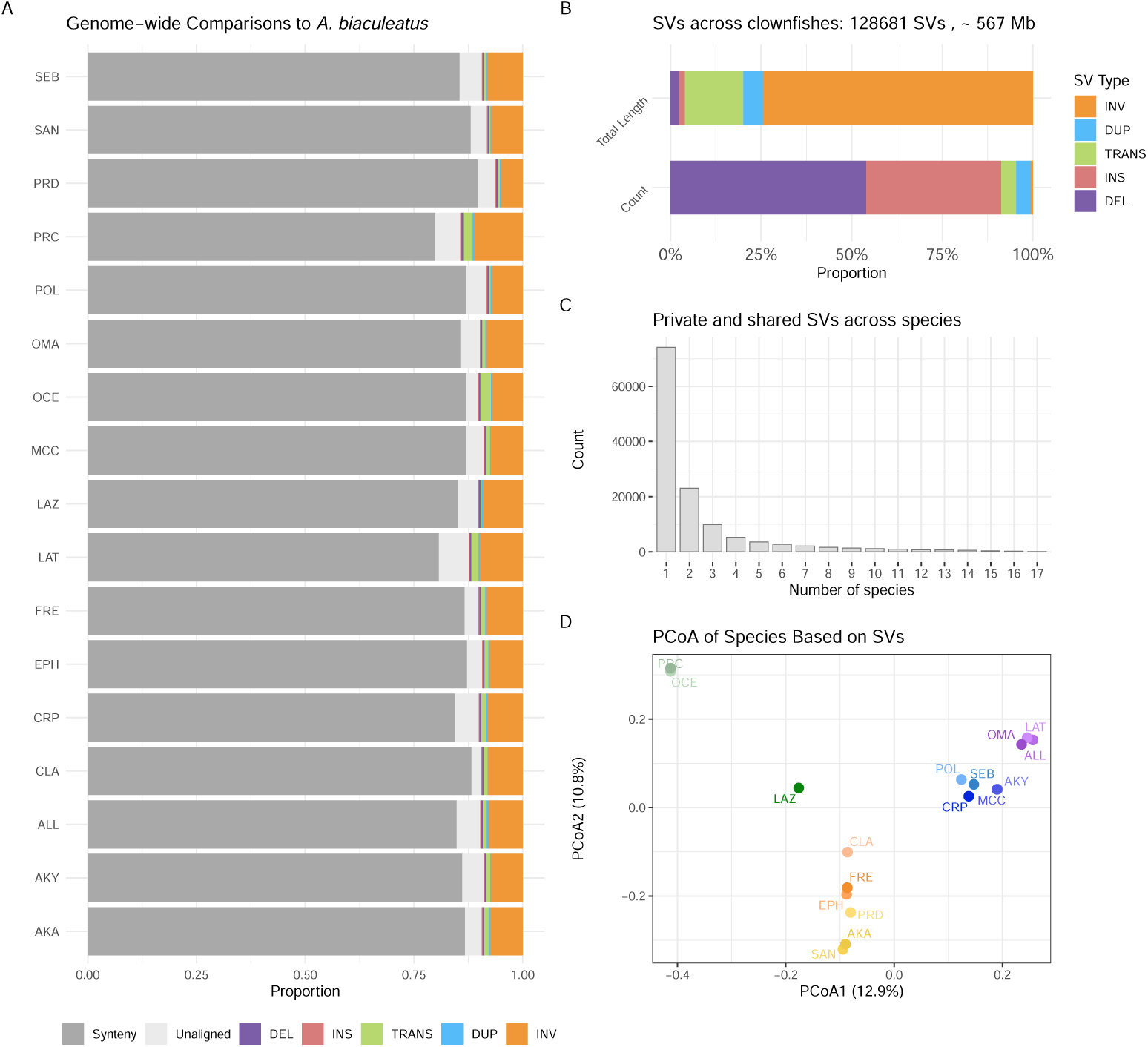
Structural variation across clownfish genomes relative to *Amphiprion biaculeatus*. Species abbreviations used throughout the figure and the study are as follows: newly assembled species are AKA = *A. akallopisos*, AKY = *A. akindynos*, ALL = *A. allardi*, CRP = *A. chrysopterus*, EPH = *A. ephippium*, FRE = *A. frenatus*, LAT = *A. latifasciatus*, LAZ = *A. latezonatus*, MCC = *A. mccullochi*, OMA = *A. omanensis*, POL = *A. polymnus*, PRC = *A. percula*, PRD = *A. perideraion*, SAN = *A. sandaracinos*, SEB = *A. sebae*; publicly available reference assemblies are CLA = *A. clarkii*, OCE = *A. ocellaris*. **A)** Proportion of each genome classified as syntenic, not aligned, or affected by a structural variant class - deletions (DEL), insertions (INS), translocations (TRANS), duplications (DUP), and inversions (INV) - relative to *A. biaculeatus*. **B)** Total count (bottom) and cumulative sequence length (top) of each SV class after merging SVs across species (i.e., SVs of the same class with start coordinates within 1 kb across species were treated as a single variant). A total of 128,681 SVs were identified, corresponding to a cumulative length of approximately 567 Mb. While INDELs are the most frequent SV class by count, inversions contribute the largest proportion of the cumulative sequence length affected. **C)** Number of SVs as a function of the number of species sharing them. The majority of SVs are private (i.e., detected in a single species), and the number of shared SVs decreases sharply as the number of species sharing them increases. **D)** Principal coordinates analysis (PCoA) based on pairwise Jaccard distances computed from SV presence/absence across all species. Point colours reflect phylogenetic groupings as inferred from the whole-genome phylogeny (Fig. 1A). Species cluster primarily according to their phylogenetic relationships rather than sea anemone host specialization. Highly similar results were obtained when performing whole-genome alignments and SV calling based on *A. clarkii* reference genome (Supplementary Figure S23).

After merging SVs across all species, we identified a total of 128, 681 SVs with a cumulative length of approximately 567 Mb (Fig. 2B). Insertions and deletions (INDELs) were by far the most abundant SV class, accounting for 91.2% of all detected variants (69, 460 deletions and 47, 890 insertions; Fig. 2B; Supplementary Figure S25A). However, these variants were generally short and collectively affected only 22.3 Mb of sequence (4.0% of the total; 13.0 Mb and 9.3 Mb for deletions and insertions, respectively; Supplementary Figure S25B-C). In contrast, inversions were less frequent (721 events, 0.6% of the detected SVs) but much larger, spanning a cumulative 422.5 Mb and representing the dominant SV class in terms of sequence length affected (74.5%; Fig. 2B; Supplementary Figure S25A–C).

The majority of SVs (57.6%) were species-specific (Fig. 2C), with *A. ocellaris* carrying the highest number (*>* 10, 000 SVs), followed by *A. percula* and *A. latezonatus* (*>* 7, 000 SVs each; Supplementary Figure S26A). The number of shared SVs decreased with the number of species sharing them (Fig. 2C), and was highest among closely related species pairs. For example, *A. ocellaris* and *A. percula* shared the most SVs (2, 962 SVs), followed by *A. akallopisos* and *A. sandaracinos* (1, 348 SVs), and by *A. akindynos* and *A. mccullochi* (1, 250 SVs). This pattern was mirrored by the principal coordinates analysis (PCoA), which clustered species primarily by phylogenetic relationships, with PCoA1 and PCoA2 explaining 12.9% and 10.8% of the variance, respectively (Fig. 2D). Similar results were obtained when considering each SV class separately (Supplementary Figure S27). Together, these results indicate that genome-wide patterns of structural variation across clownfish species primarily reflect shared ancestry rather than ecological factors. However, because PCoA summarizes overall genomic patterns, it cannot determine whether particular subsets of SVs are associated with specialization to similar sea anemone hosts. We therefore explicitly tested for associations between individual structural variants and host specialization.

### Structural variation and host specialization

We identified candidate host-specific SVs as variants present in all species within a given host category and absent from all other species. Only a small number of such SVs were detected in *Radianthus* specialists (*n* = 12, mostly deletions) and *Stichodactyla* specialists (*n* = 59, mostly insertions), whereas no SVs were shared across all *Entacmaea* specialists (Supplementary Figure S28; Supplementary Table S10). Of the 71 host-specific SVs identified, only seven overlapped protein-coding sequences and could potentially affect gene function (Supplementary Table S10). However, none of these genes showed a clear functional link to clownfish ecology or host specialization.

To assess whether these shared SVs were more common than expected by chance, we compared their number to the number of SVs shared among equally sized control groups containing species with different host associations. We found no excess of shared SVs in either specialist group (Supplementary Figure S28). In *Stichodactyla* specialists, the 59 SVs shared by *A. polymnus* and *A. sandaracinos* fell within the range observed for the three control pairs (53–71 SVs; Supplementary Figure S28). Likewise, the 12 SVs shared among the four *Radianthus* specialists were similar to the 10 SVs shared by the control group (Supplementary Figure S28). Together, these results suggest that SVs shared among host specialists are likely due to chance rather than adaptation to particular sea anemone hosts. Consistent results were obtained when analyses were performed using *A. clarkii* as the reference genome (Moore et al., 2023) (Supplementary Figure S4).

### Large inversions and topological inconsistencies

Although no SV was clearly associated with host specialization, structural variation may still contribute to clownfish diversification more broadly. To support future work in this direction, we compiled a catalogue of all SVs identified in this study, distinguishing private and shared variants and annotating those overlapping protein-coding sequences (CDS; SV Catalogue). Interestingly, when only considering SVs spanning CDS, the total number of SVs dropped to 10,077, with 6,452 SVs being species-specific and 3,625 being shared across species, indicating that most SVs do not directly affect protein-coding sequences.

Among the different SV classes catalogued, we focused on large inversions, since they are more likely to suppress recombination between the standard and inverted arrangements, potentially maintaining linked sets of variants and contributing to adaptation (Mérot et al., 2020; Wellenreuther et al., 2025). Because recombination suppression potentially produces local phylogenies that differ from the genome-wide species tree, we screened the SV catalogue for long inversions and examined whether they were associated with topological inconsistencies relative to the genome-wide phylogeny.

Out of the 110 large (*>* 100 kb) inversions shared by at least two species, 68 were located in genomic regions with sequence alignments available for at least nine species. Although topological discordance was detected within the inversions, similar inconsistencies were observed across the genome and were not clearly associated with the presence or absence of inversions (Supplementary Figure S29). Two exceptions were nevertheless observed: an inversion on chromosome CM132566.1 (homologous to chromosome 18 in *A. clarkii* reference genome) and an inversion on chromosome CM132553.1 (homologous to chromosome 9 in *A. clarkii* reference genome; Fig. 3).

**Figure 3:**
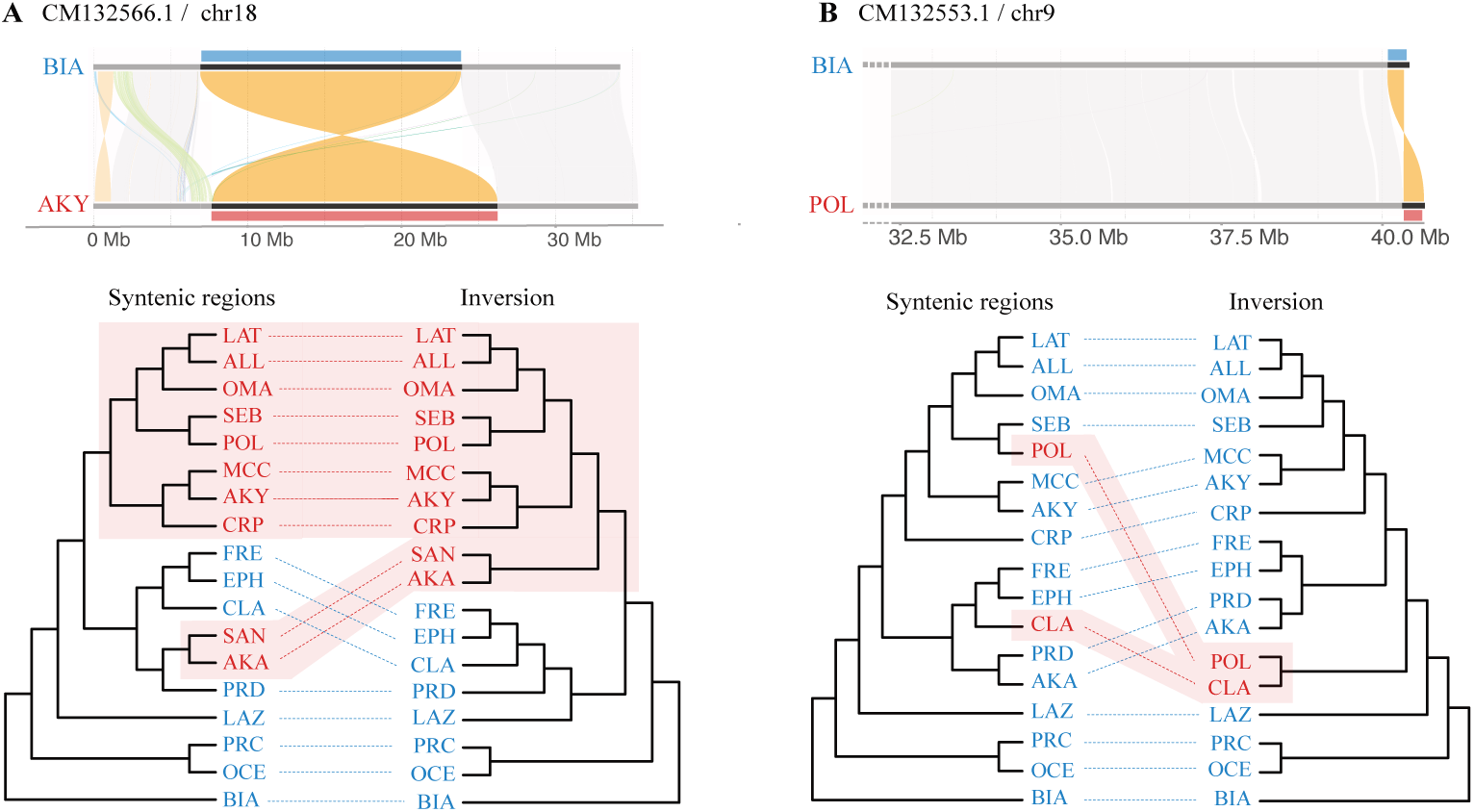
Discordant local topologies associated with long inversions on chromosomes CM132566.1/chr18 (**A**) and CM132553.1/chr9 (**B**). Chromosome names refer to the *A. biaculetus* reference genome (CM132566.1, CM132553.1) and the homologous chromosomes in the *A. clarkii* reference genome (chr18, chr9). For both loci, the top panel shows an example alignment between *A. biaculeatus* (BIA) and a representative species carrying the inverted haplotype, with syntenic regions, translocations, and inversions shown in grey, green, and orange, respectively; the “standard” and “inverted” arrangements are shown in blue and red, respectively, with the standard arrangement defined as that found in *A. biaculeatus*, the basal species in the genus. Similar alignments were observed across all species carrying either arrangement. The bottom panel shows cladograms of the phylogenetic relationships inferred from SNPs extracted from the syntenic regions flanking the inversion (left) and from SNPs extracted from the inverted region itself (right); species carrying the inverted haplotype are highlighted in red, and species with the standard haplotype are shown in blue. In both cases, the inverted-region topology shows clear discordance, with species carrying the inversion recovered as a monophyletic group. **A)** Alignment shown for *A. akindynos* (AKY) relative to BIA. **B)** Alignment shown for *A. polymnus* (POL) relative to BIA

The inversion on CM132566.1 (chr18) spanned approximately 17 Mb and was shared by 10 species (*Amphiprion akallopisos*, *A. sandaracinos*, *A. chrysopterus*, *A. mccullochi*, *A. akindynos*, *A. sebae*, *A. polymnus*, *A. omanensis*, *A. allardi*, and *A. latifasciatus*; Fig. 3A). Phylogenetic reconstruction based on SNPs within the inversion recovered all inversion-carrying species as a monophyletic group, with the two sister species *A. akallopisos* - *A. sandaracinos* moving to be basal to the others instead of being placed as sister to *A. perideraion* (Fig. 3A). Analysis of inversion breakpoints revealed no evidence of heterozygosity in any of the species carrying the inversion (Supplementary Table S11A). Furthermore, no repetitive elements were detected directly at the breakpoints, and no significant enrichment or depletion of repeat elements was observed at breakpoint regions relative to the rest of the genome (Supplementary Table S12A).

In most species (*A. akallopisos*, *A. akindynos*, *A. allardi*, *A. chrysopterus*, *A. mccullochi*, *A. sandaracinos*, and *A. sebae*), the distal breakpoint was located within 5 kb of two genes: *myct1b* (myc target 1b), located downstream of the breakpoint, and *tulp4b* (TUB-like protein 4b), located upstream. At the proximal breakpoint, *alkbh1* (AlkB homolog 1, histone H2A dioxygenase) was located within 5 kb downstream of the breakpoint. Depending on the species, the inversion encompassed between 611 and 717 annotated genes, including the growth and appetite-related gene *ghsra* (ghrelin receptor) and *ltk*, a receptor tyrosine kinase repeatedly implicated in iridophore and pigment-pattern evolution in teleosts. The inverted region also contained several developmental transcription factors associated with eye and forebrain/thyroid patterning, including *six1*, *six4a*, *six6*, *vsx2*, *pax1a*, *pax9*, *foxa2*, *sox11a*, and members of the *nkx2* homeobox family (Supplementary Table S13). Gene Ontology (GO) enrichment analysis identified 18 significantly enriched terms (Supplementary Table S14A), including terms related to animal organ morphogenesis (GO:0009887), cerebral cortex development (GO:0021987), and regulation of behavior (GO:0050795). Notably, the inversion contained the serotonin receptor genes *htr1b* and *htr1d* (Supplementary Table S13).

The inversion on CM132553.1 (chr9) was identified only in *A. polymnus* and *A. clarkii*. Phylogenetic reconstruction based on SNPs within this 256 kb inversion recovered these two species as sister taxa (Fig. 3B). As in the CM132566.1/chr18 inversion, breakpoint analyses found no evidence of heterozygosity (Supplementary Table S11B), and no significant enrichment or depletion of repetitive elements was detected at the breakpoint regions (Supplementary Table S12B). In both species, the *cullin-5-like* gene was located approximately 2.5 kb upstream of the distal breakpoint, whereas no genes were found near the proximal breakpoint. The inversion contained 15 annotated genes in both species (Supplementary Table S15), including *capn5a*, encoding the calcium-dependent cysteine protease calpain 5a, and the gene *ompa*, coding for the olfactory marker protein. GO enrichment analysis identified a single significantly enriched term shared by both species, GO:0032652 (regulation of interleukin-1 production) (Supplementary Table S14B).

## Discussion

In this study, we generated high-quality genome assemblies for 16 clownfish species, enabling the first genome-wide characterization of structural variation across the clade. We identified nearly 130,000 SVs spanning close to 570 Mb; short indels were by far the most numerous class, but large inversions accounted for most of the sequence affected. While we found no clear association between structural variation and sea anemone host specialization, we resolved a single large inversion of approximately 17 Mb on chromosome 18, clarifying earlier reports of two separate rearrangements at this locus (Schmid et al., 2022; Marcionetti and Salamin, 2023; Marcionetti et al., 2024), and identified a second, smaller inversion on chromosome 9. Both inversions show evolutionary histories distinct from the rest of the genome, consistent with a history of hybridization.

All 16 assemblies obtained in this study achieved Quality Values (QV) above 43, exceeding values reported in comparable long-read assembly studies (Krawczyk et al., 2025; Maughan et al., 2024; Parey et al., 2024; Pu et al., 2024). Combined with high BUSCO (Simão et al., 2015) and OMArk (Nevers et al., 2025) completeness and low structural and small-scale error rates, these genomes represent an important new resource, bringing the number of high-quality clownfish assemblies available to the community up to 18 of the 28 currently recognized species. Nevertheless, despite their consistently high base-level accuracy and completeness, it is important to notice that these are not telomere-to-telomere assemblies. No Hi-C or other long-range scaffolding data were generated for this study, which are generally required to fully resolve the remaining gaps and repetitive regions in chromosome-scale assemblies (Li and Durbin, 2024). Instead, scaffolding relied exclusively on HiFi-read-based approaches. Consequently, a few scaffolds remain unplaced or unoriented relative to a complete chromosome-scale karyotype, and some fragmentation persists despite successive rounds of scaffolding (Fig. 1A; Supplementary Table S4). Nevertheless, the overall accuracy, completeness, and contiguity of these assemblies are consistently high, making them well suited for the comparative analyses performed here and providing a robust foundation for future genomic studies as taxonomic sampling across the genus continues to expand.

The newly assembled reference genomes allowed a first broad quantification of structural variation across clownfishes. SVs were found to affect between 6.3% and 14.3% of the genome when each species was compared with *A. biaculeatus*, at the base of the clownfish phylogeny. This proportion was higher than the pairwise single-nucleotide divergence found across the same comparisons, which ranged between 1.51% to 1.61% and indicating that, when considering SVs, clownfish genomes are between 4 and 10 times more different than when considering SNPs alone. This result is consistent with recent comparative work across vertebrates, where authors analysed over 600 species and 500 million years of evolution and found that SVs were found to impact roughly 12-fold more base pairs than SNPs genome-wide (Lou et al., 2026). A similar pattern has also been reported within fish radiations. Indeed, in Lake Malawi cichlids, pairwise SNP divergence is remarkably low (0.1–0.25%) yet structural variation accounts for close to 5–10% of the genome, which suggests that SVs represent a source of genomic diversity that SNP-based studies alone systematically underestimate (Quah et al., 2025). Our results in clownfishes fit this broader pattern, reinforcing the view that structural variants capture a larger component of genomic divergence between species than SNPs alone (Mahmoud et al., 2019) and underscoring the importance of incorporating them into comparative genomic analyses.

INDELs were by far the most abundant class of SVs, whereas inversions, despite being much rarer, accounted for the largest proportion of affected genome length. This pattern was consistent both in pairwise comparisons of each species against *A. biaculeatus* and in the merged dataset across species. It was also recovered when *A. clarkii* was used as the reference (Supplementary Figure S23, Supplementary Table S9), indicating that it is highly unlikely to be an artifact of genome reference choice. In addition, the same asymmetry was observed within individuals when considering heterozygous SVs identified within each assembly (Supplementary Figure S22).

The much larger number of INDELs relative to inversions is expected: multiple INDELs can occur within the span of a single large inversion. However, this counting effect alone is unlikely to explain the magnitude of the observed difference. Similar patterns have been reported in studies using different detection approaches and spanning diverse taxa. For example, a Lake Malawi cichlid pangenome identified more than 100,000 insertions and 100,000 deletions but fewer than 1,000 inversions (Quah et al., 2025), while a comparable excess of INDELs over inversions has also been reported in *Theobroma cacao* (Hämälä et al., 2021). This imbalance likely reflects differences in both the mutational mechanisms generating each SV class and the selective forces acting upon them. INDELs can arise through mechanisms involving short regions of microhomology, replication errors, or small-scale repair events, whereas inversions typically require non-allelic homologous recombination between long, highly similar repeats in opposite orientations at both breakpoints, a substantially rarer genomic configuration. Consequently, inversions are expected to occur less frequently than INDELs (Carvalho and Lupski, 2016; Burssed et al., 2022). In addition, theoretical and simulation studies predict that inversions experience stronger purifying selection on average because suppressed recombination between inverted and standard arrangements can promote the accumulation of deleterious mutations within inverted haplotypes, increasing their genetic load (Berdan et al., 2021a).

Although inversions can reduce fitness by allowing deleterious mutations to accumulate, they are also frequently implicated in local adaptation. The recombination suppression that results in the accumulation of deleterious alleles can also preserve co-adapted combinations of locally beneficial alleles (Kirkpatrick and Barton, 2006; Faria et al., 2019; Wellenreuther et al., 2025). Beyond inversions, other forms of structural variation have also been linked to adaptation across taxa. For example, CNVs contribute to plumage colouration and thermal adaptation in the common murre (Tigano et al., 2018). Moreover, the same SV can sometimes contribute to recurrent adaptation across multiple species within a radiation, either through repeated introgression or through the repeated recruitment of ancestral polymorphism. Examples of such patterns have been described in the Lake Malawi cichlids (Blumer et al., 2025) and in *Lonchura* (Knief et al., 2024) radiations, where the same inversions have been implicated in recurrent ecological adaptation across multiple species. We thus asked whether a similar pattern of repeated use of the same structural variants could be detected in clownfishes diversification.

Overall, most SVs were species-specific, while shared SVs largely reflected phylogenetic relationship, suggesting that structural variation across clownfishes primarily reflects lineage history rather than repeated ecological adaptation. Focusing specifically on host use, host-specific variants were rare and occurred no more frequently than expected by chance (Supplementary Figures S28–S4), and the few coding sequences they overlapped had no obvious functional link to host specialization (Supplementary Table S10). Thus, recurrent specialization on the same sea anemone hosts does not appear to be associated with repeated presence of the same SVs across species. We note that our analysis specifically tested for SVs shared among all species associated with a given sea anemone host. It therefore remains possible that ecologically relevant SVs are shared among only a subset of host-associated species and would not be detected here. Nevertheless, convergence associated with host specialization appears to be primarily associated with convergent positive selection at particular protein-coding genes (Gaboriau et al., 2025), rather than by recurrent changes in structural variation.

While mostly reflecting phylogenetic relationship, we nonetheless found SVs showing a paraphyletic distribution across species, i.e., present in some but not all species within a clade (see SV Catalogue). Assembly fragmentation could partially explain this pattern, as it may have prevented the detection of some SVs in certain species, thus resulting in their apparent absence. Nevertheless, this pattern could also be biological rather than technical. The same SVs may have arisen independently in different species or populations and may have been fixed through natural selection (e.g., Cáceres et al., 2007; Porubsky et al., 2020; Pokrovac et al., 2024). Alternatively, SVs may have been present as ancestral standing genetic variation and subsequently fixed in only a subset of descendant species, or transferred between species through interspecific hybridization and introgression (e.g., Jones et al., 2012; Jay et al., 2018; Knief et al., 2024; Blumer et al., 2025), resulting in the paraphyletic distribution pattern sometimes observed here. Finally, because our analysis includes only one individual per species, some of the observed SVs may represent within-species polymorphisms rather than fixed differences between species, and the observed paraphyletic pattern could therefore partly reflect incomplete sampling of intraspecific variation. Distinguishing among these alternative scenarios, and determining whether any of these SVs have functional consequences, will require more comprehensive within-species sampling alongside additional population-level and comparative analyses. Thus, while these para-phyletically distributed SVs may provide interesting candidates for further investigation, their evolutionary history and potential functional significance cannot currently be inferred with confidence.

A partial exception to this is the case of inversions showing clear topological inconsistency, where species carrying the inversion branch together and appear more closely related within the inversion than they do genome-wide. Two large inversions showing this pattern were previously hypothesized on chromosome 18 (relative to the *A. clarkii* and *A. percula* reference genomes; homologous to CM132566.1 in the *A. biaculeatus* reference genome) (Schmid et al., 2022; Marcionetti and Salamin, 2023; Marcionetti et al., 2024). We confirmed here the presence of a single large inversion of approximately 17 Mb on this chromosome (Fig. 3A). The two inversions hypothesized in previous studies likely reflect an artifact of the *A. percula* reference genome used (GCA 003047355.2), which appears to carry an independent inversion at the end of chromosome 18 (Supplementary Figure S2), splitting the true inversion into two blocks. This independent inversion was not recovered in the *A. percula* assembly generated here, and is therefore not discussed further.

When looking in more detail at the CM132566.1/chr18 inversion, the monophyly of the carrier species, together with the presence of the same breakpoints across species, suggests that it originated as a single event. This hypothesis is further supported by the low repeat content observed at the breakpoints, in contrast to the elevated repeat density typically associated with regions of high breakpoint turnover and recurrent rearrangement (Gozashti et al., 2025). Under this scenario, the inversion arose once from an ancestral non-inverted haplotype (as defined by the basal species of the group, *A. biaculeatus*) in the ancestor of the Indian Ocean/*polymnus*/Australian/*chrysopterus* clade (following the clade definitions of Litsios and Salamin, 2014). The inversion may then have spread in the ancestor of *A. akallopisos* and *A. sandaracinos* through ancestral hybridization. At freely recombining loci, introgression signals are typically eroded by recombination over time; within inversions, by contrast, the absence of recombination with the alternative non-inverted haplotype allow signatures of ancestral hybridization events to persist for much longer (Sturtevant and Beadle, 1936).

While we did not formally test for this here, introgressive hybridization on chromosome 18 has been previously demonstrated (Schmid et al., 2022; Marcionetti and Salamin, 2023), favoring hybridization over incomplete lineage sorting as the more likely explanation for the topological signal observed at this locus. *A. chrysopterus* and *A. sandaracinos* still hybridize today, producing the naturally occurring hybrid *A. leucokranos* in Papua New Guinea and the Solomon Islands (Gainsford et al., 2015, 2020; Schmid et al., 2025). This shows that reproductive isolation between these two lineages remains incomplete, making similar but older hybridization events a plausible mechanism for the origin and spread of the CM132566.1/chr18 inversion.

When looking at the potential function of the CM132566.1/chr18 inversion, we see that it contains several genes potentially involved in development and behaviour. Among them, we found a receptor tyrosine kinase *ltk* that is required for the specification, proliferation, and survival of iridophores in zebrafish (Lopes et al., 2008; Fadeev et al., 2016; Mo et al., 2017), the latter being the cell type composing the white bars in clownfish species (Salis et al., 2019). The inversion also contains a cluster of transcription factors patterning the eye and the forebrain/thyroid axis, including *six1*, *six4a*, *six6*, *vsx2*, *pax1a*, *pax9*, *foxa2*, *sox11a*, and members of the *nkx2* homeobox family, genes whose vertebrate orthologs govern retinal progenitor identity and thyroid organogenesis (Zuber et al., 2003; Diacou et al., 2018; Ferrena et al., 2024). This combination is interesting given that clownfish thyroid hormones regulate the formation of white bars in clownfishes (Salis et al., 2021) and that larval-to-juvenile metamorphosis is itself a thyroid-hormone-controlled transition that reorganizes the visual system, including a shift in opsin expression as larvae move from the pelagic environment to settle on the reef (Roux et al., 2023). The inversion also contains the serotonin receptor genes *htr1b* and *htr1d*, previously linked to dominance behavior in fishes (Trainor and Hofmann, 2006). Taken together, the presence of pigmentation, sensory, developmental, and behavioral genes within a single, non-recombining block could point toward the inversion acting as a supergene coordinating multiple, potentially co-adapted traits (Kirkpatrick and Barton, 2006; Faria et al., 2019; Wellenreuther et al., 2025). Nevertheless, no phenotype currently known to us unites *A. akallopisos* and *A. sandaracinos* with the other eight carrier species, nor is the inversion linked to host specialization. Formally testing the functional importance of this inversion and its genes, their potential role in adaptation, and the role of selection in fixing them into species or populations will require further studies beyond the analysis presented here.

In this study we also identified a second, smaller (approximately 256 kb) inversion on chromosome CM132553.1 (in the *A. biaculeatus* reference genome; homologous to chromosome 9 in *A. clarkii*), showing a monophyletic relationship between the two species carrying it, *A. polymnus* and *A. clarkii* (Fig. 3B). These are the two species considered the most phenotypically polymorphic among clownfishes (Salis et al., 2018; Schmid et al., 2024). The estimated divergence time between the two species (around 6 Mya; Litsios et al., 2012; Gaboriau et al., 2025) suggests that hybridization, rather than incomplete lineage sorting, is the more likely explanation for this pattern. As with the CM132566.1/chr18 inversion, the absence of recombination within the inversion may have allowed a signature of such introgressive hybridization to persist, even as it was erased elsewhere in the genome. This scenario is also plausible given that the two species co-occur around the Coral Triangle region (although recent evidence suggests that both *A. polymnus* and *A. clarkii* could be species complexes; Fitzgerald et al., 2026a, Zwahlen et al., in prep). *A. clarkii* is also the species with the widest geographic range and the most generalist host use, occurring across all ten sea anemone host species (Litsios et al., 2012; Gaboriau et al., 2025). However, contrary to chr18, this inversion contains no genes with an obvious functional link to phenotypes, and further studies would be needed to determine whether the inversion is fixed in species and populations, and whether it carries any adaptive significance.

This study provides the first genome-wide characterization of structural variation across the clownfish radiation, establishing the extent and diversity of SVs relative to SNP divergence and identifying candidate regions, including the chr18 and chr9 inversions, for further genomic investigation. However, several questions remain open. Because our dataset relies on a single genome per species, we cannot currently distinguish SVs that are fixed within species from those that remain polymorphic, a distinction that will require population-level genomic data to resolve. A pangenomic approach incorporating multiple individuals per species would allow both within- and between-species structural variation to be quantified more precisely, and represents a natural next step for this system. Timing the origin of specific SVs, and formally testing among the alternative evolutionary scenarios that could explain their present-day distribution, will likewise require dedicated simulation-based approaches; such analyses, however, are only meaningful once a more complete picture of within- and between-species SV variation is available. In providing high-quality chromosome-level genomic resources for 16 clownfish species alongside the first characterization of their structural variation, this study takes an important step toward understanding the extent and evolutionary significance of structural variation in this iconic radiation, while laying the groundwork for the population-level and simulation-based work needed to resolve the questions it raises.

## Acknowledgements

We thank Rosanna Pescini Gobert for lab management. We also thank the Lausanne Genomic Technologies Facility for sequencing the samples and the DCSR infrastructure of the University of Lausanne for the computational resources. We thank Nicolas Rodrigues for the clownfish drawings. The work was funded by a grant from the Swiss NSF to N.S. (grant 310030 185223) and by funding from the University of Lausanne.

## Data Accessibility Statement

Scripts are available on Github (https://github.com/amarcion/ClownfishSV/). The SV catalogue is available at https://amarcion.github.io/ClownfishSV/ClownfishSV_database.html. Data (raw reads, genome assemblies, genome annotations) are deposited in the European Nucleotide Archive (ENA; Study ID: PRJEB126889) and will be made publicly available upon acceptance of the manuscript.

## Benefits-Sharing Statement

The sharing of our data and results on public databases ensures transparency and facilitates further research in the field. All fieldwork conducted during this study adhered to local regulations and was carried out in collaboration with local entities. Our research activities were conducted in accordance with the “Access and Benefit Sharing” (ABS) principles of the Nagoya Protocol established by the Convention on Biological Diversity (CBD).

## Author Contributions

AM and NS conceptualised and designed the study. AM conducted the research, performed the analyses, interpreted the results and wrote the manuscript. EIM contributed to genomic assembly, annotation and structural variant identification. DAH, SS and NS contributed to data interpretation. DAH contributed to the manuscript writing. SS contributed to the inversion analyses. Fieldwork expeditions and sample collection were carried out by FC, LMF, BF, TG, AGJ, VL. All authors reviewed, revised and approved the final version of the manuscript.

