## Supplementary Figure for "Genomic structural variation in clownfish adaptive radiation"

### Supplementary materials

#### Contents

**1 Supplementary Table Legends** **S1**

**2 Supplementary Figures** **S5**

### Supplementary Tables

#### 1 Supplementary Table Legends

**Table S1. Information on the samples used in the study.** For each of the 16 species, the table reports the species name, abbreviation used throughout the study, original and additional sample identifiers, sample origin, sampling location, sequencing date, resequencing date where applicable, and sequencing technology. Samples were obtained through fieldwork expeditions, collaborators at the Marine Station of Academia Sinica (Taiwan), or aquarium suppliers.

**Table S2. Assembly statistics obtained with four different assemblers evaluated on four species.** To select the most suitable assembler for the dataset, four software packages were tested on *A. akallopisos*, *A. ephippium*, *A. perideraion*, and *A. polymnus*: IPA (v1.8.0), Flye (v2.9.1), HiCanu (v2.2), and Hifiasm (v0.19.5). For HiCanu assemblies, haplotig duplications were removed with purge_dups (v1.2.5) prior to evaluation. The table reports for each species–assembler combination: total assembly size (Mbp), number of contigs, mean contig length (kbp), longest contig (Mbp), shortest contig (bp), gap count (Ns), N50 and N90 values and their corresponding counts, and BUSCO completeness (complete %, complete single-copy %, and missing %; *actinopterygii_odb10* database). Hifiasm consistently produced the most complete and contiguous assemblies and was selected for all species.

**Table S3. Assembly statistics and quality metrics at each curation step for all 16 species.** For each species and assembly step, the table reports: number of contigs, number of contigs longer than 1 Mbp, total assembly length (bp), total length of contigs >1 Mbp, longest and second-longest contig (bp), N50 (Mb), HiFi read mapping rate (%), split-read rate (%), sequencing depth, mapping rate in large contigs (%), Quality Value (QV), and counts of structural errors (expansions, collapses, haplotype switches, inversions), small-scale assembly errors per Mbp, total small-scale errors, base substitutions, small-scale expansions, and small-scale collapses. All metrics were computed with Inspector (v1.3.1). Assembly steps follow the same numbering as Supplementary Figure S1.

**Table S4. Overall assembly statistics, quality and completeness for the final assemblies of all 16 species**. For each species, the table reports: **A)** assembly statistics including total assembly length (Gbp), number of scaffolds, mean and longest scaffold length (Mbp), shortest scaffold length (bp), total gap count and gap length (Ns), N50 and N90 scaffold lengths (Mb) and their corresponding counts

(N50n, N90n); **B)** assembly quality estimated with Inspector (v1.3.1), reported as Quality Value (QV); and **C)** assembly completeness estimated with BUSCO (v5.4.7; *actinopterygii_odb10* database; 3,640 genes), reporting the percentage of complete, single-copy, duplicated, fragmented, and missing BUSCOs.

**Table S5. Sequencing statistics for all 16 species**. For each species, the table reports the sample identifier, total number of HiFi reads, sequencing yield (Gbp), mean read length (bp), and median read length (bp). Reads were generated on PacBio Sequel II or Revio platforms using Circular Consensus Sequencing (CCS).

**Table S6. Repeat content for all 16 species.** For each species, the table reports the total assembly length (Mbp), GC content, total masked sequence (Mbp and %), and the count and percentage of the genome occupied by each repeat class: SINEs, LINEs, LTR elements, DNA transposons, unclassified repeats, total interspersed repeats, small RNAs, satellites, simple repeats, and low-complexity regions. Repeat libraries were modelled with EDTA (v2.1.3) and RepeatModeler2 (v2.0.1), and assemblies were masked with RepeatMasker (v4.0.7).

**Table S7. Genome annotation statistics for all 16 species.** For each species, the table summarizes gene annotation statistics, including the total number of predicted protein-coding genes, the proportion and number of functionally annotated genes, and the total number of transcripts. Annotation completeness as assessed by BUSCO (v5.4.7; actinopterygii_odb10 database; 3,640 genes) and by OMArk (v0.3.1; Metazoa database; 16,037 HOGs) is also reported, with the percentage of complete (single-copy and duplicated), fragmented, and missing genes.

**Table S8. Genomic comparison statistics for all species relative to the *A. biaculeatus* reference genome.** For each species, the table reports the total analysed genome length (bp) and summarizes both counts and cumulative lengths (bp) of syntenic regions, unaligned regions, and structural variant classes (inversions, translocations, duplications, and insertions and deletions larger than 50 bp). The percentage of the genome affected by structural variants, the total number of SNPs within syntenic regions, and SNP divergence in % are also reported. Structural variants were identified using SyRI (v1.6.3) from whole-genome alignments generated with minimap2 (v2.28; -x asm5).

**Table S9. Genomic comparison statistics for all species relative to the *A. clarkii* reference genome.** For each species, the table reports the total analysed genome length (bp) and summarizes both counts and cumulative lengths (bp) of syntenic regions, unaligned regions, and structural variant classes (inversions, translocations, duplications, and insertions and deletions larger than 50 bp). The percentage of the genome affected by structural variants, the total number of SNPs within syntenic regions, and SNP divergence in % are also reported. Structural variants were identified using SyRI (v1.6.3) from whole-genome alignments generated with minimap2 (v2.28; -x asm5).

**Table S10. Host-specific structural variants (SVs), relative to *A. biaculeatus*.** Full list of the 71 SVs shared by all species within a given sea anemone host category (*Radianthus*, $n = 12$; *Stichodactyla*, $n = 59$) and absent from all other species. For each SV, the table reports the chromosome and position, length, SV identifier and type, the species sharing it, the host category, whether it overlaps a protein-coding sequence (CDS), and, for the seven SVs overlapping CDS, the affected transcript identifier and the corresponding best DIAMOND BLASTP hit (NR database) with protein name.

**Table S11. Continuous and split reads at inversion breakpoints.** For each of the two breakpoints of the chromosome 18 (**A**) and chromosome 9 (**B**) inversions, the table reports, for every carrier species, the number of HiFi reads mapped back to the corresponding assembly across the breakpoint, the number of split (SA-tagged) reads, the number of continuous reads, and the resulting split-read ratio. Continuous reads support the haplotype present in the individual's own assembly, whereas split reads spanning the breakpoint would indicate the presence of the alternative arrangement. Ratios close to zero across all carrier species indicate no evidence of heterozygosity at either breakpoint.

**Table S12. Repeat content at inversion breakpoints.** For each breakpoint of the chromosome 18 (**A**) and chromosome 9 (**B**) inversions, the table reports, for every carrier species, the amount and proportion of repetitive sequence within a window centred on the breakpoint, alongside the mean, standard deviation, and number of equivalently sized background windows sampled across the rest of the genome. An empirical two-tailed permutation p-value tests whether breakpoint windows are significantly enriched for repetitive sequence relative to this genomic background.

**Table S13. Genes located within the CM132566.1/chr18 inversion.** List of genes falling within the boundaries of the ∼17 Mb inversion on chromosome CM132566.1/chr18, grouped as Hierarchical Orthologous Groups (HOGs) inferred across the 10 carrier species (*A. akallopisos*, *A. akindynos*, *A. allardi*, *A. chrysopterus*, *A. latifasciatus*, *A. mccullochi*, *A. omanensis*, *A. polymnus*, *A. sandaracinos*, *A. sebae*). For each HOG, the table reports the HOG identifier, the number of species in which it was recovered, gene name and description, and the corresponding per-species gene identifier.

**Table S14. Gene Ontology (GO) enrichment among genes within the chromosome 18 (A) and chromosome 9 (B) inversions.** GO enrichment was tested independently within each carrier species using topGO (weight01 algorithm), considering genes located within the inversion boundaries against a genome-wide background. The table lists GO terms significant in every carrier species tested (10 of 10 species for the chromosome 18 inversion; 2 of 2 species, *A. polymnus* and *A. clarkii*, for the chromosome 9 inversion), reporting the GO identifier, term description, number of species in which the term was significant out of the number tested, and the per-species enrichment p-value. No correction for multiple testing across GO terms or species was applied.

**Table S15. Genes located within the CM132553.1/chr9 inversion.** List of the 15 genes falling within the boundaries of the ∼256 kb inversion on chromosome CM132553.1/chr9, shared by *A. polymnus* and *A. clarkii*. For each Hierarchical Orthologous Group (HOG), the table reports the HOG identifier, the number of species in which it was recovered, gene name and description, and the corresponding gene identifier in each of the two carrier species.

### 2 Supplementary Figures

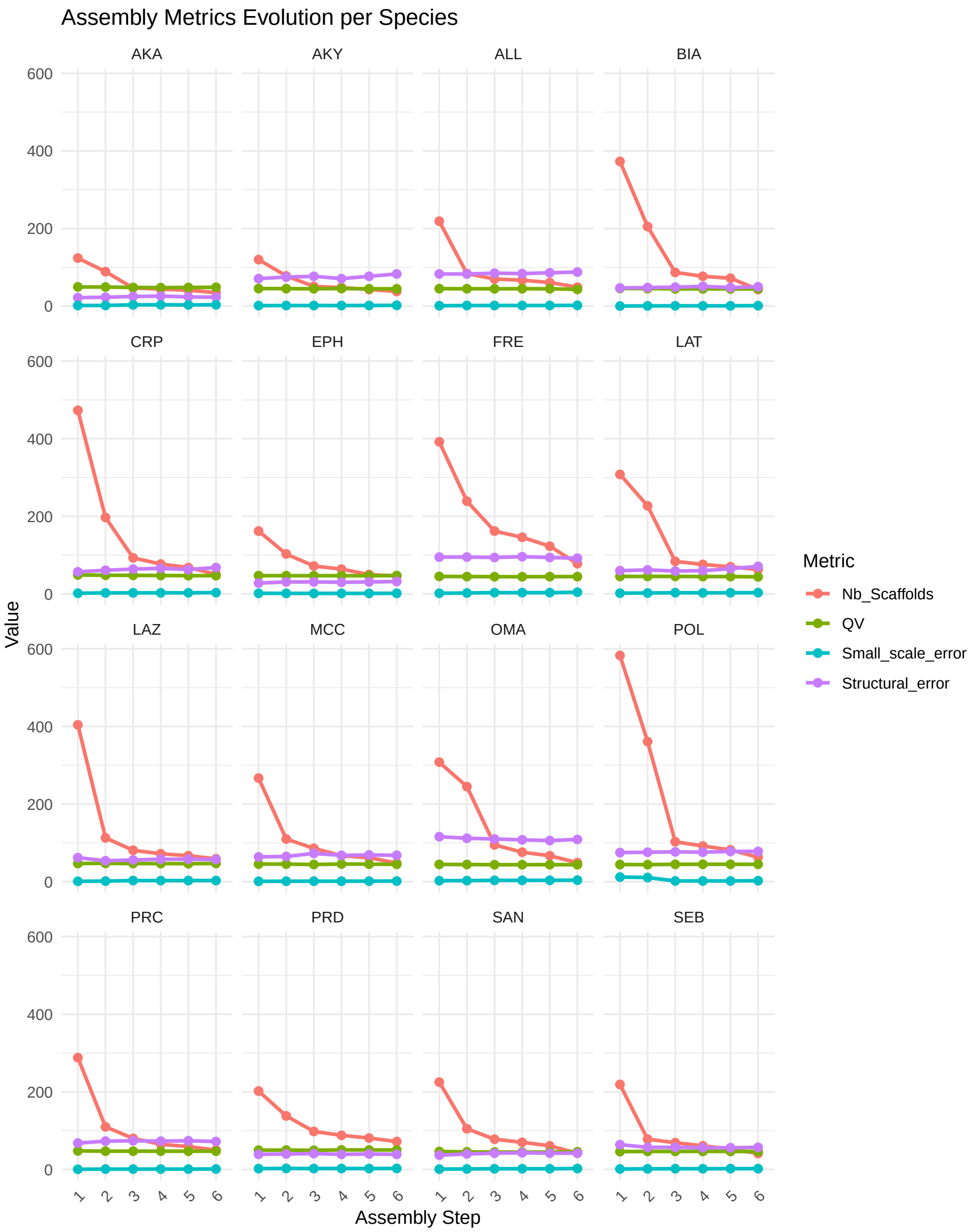


Figure S1: Evolution of assembly metrics across the successive curation steps for all 16 species. For each species (facet), four metrics are tracked: number of scaffolds, Quality Value (QV), number of small-scale errors per Mbp, and number of structural errors. QV and error counts were computed with Inspector (v1.3.1). Assembly steps are: 1: initial Hifiasm assembly; 2: removal of haplotig duplications with purge_dups; 3: removal of mitochondrial and duplicated sequences; 4: scaffolding with SAMBA; 5: scaffolding with ntLink; 6: final assembly after scaffolding with P_RNA_scaffolder and removal of scaffolds shorter than 200 kb. Each step reduced scaffold number while maintaining high QV and low error rates across all species.

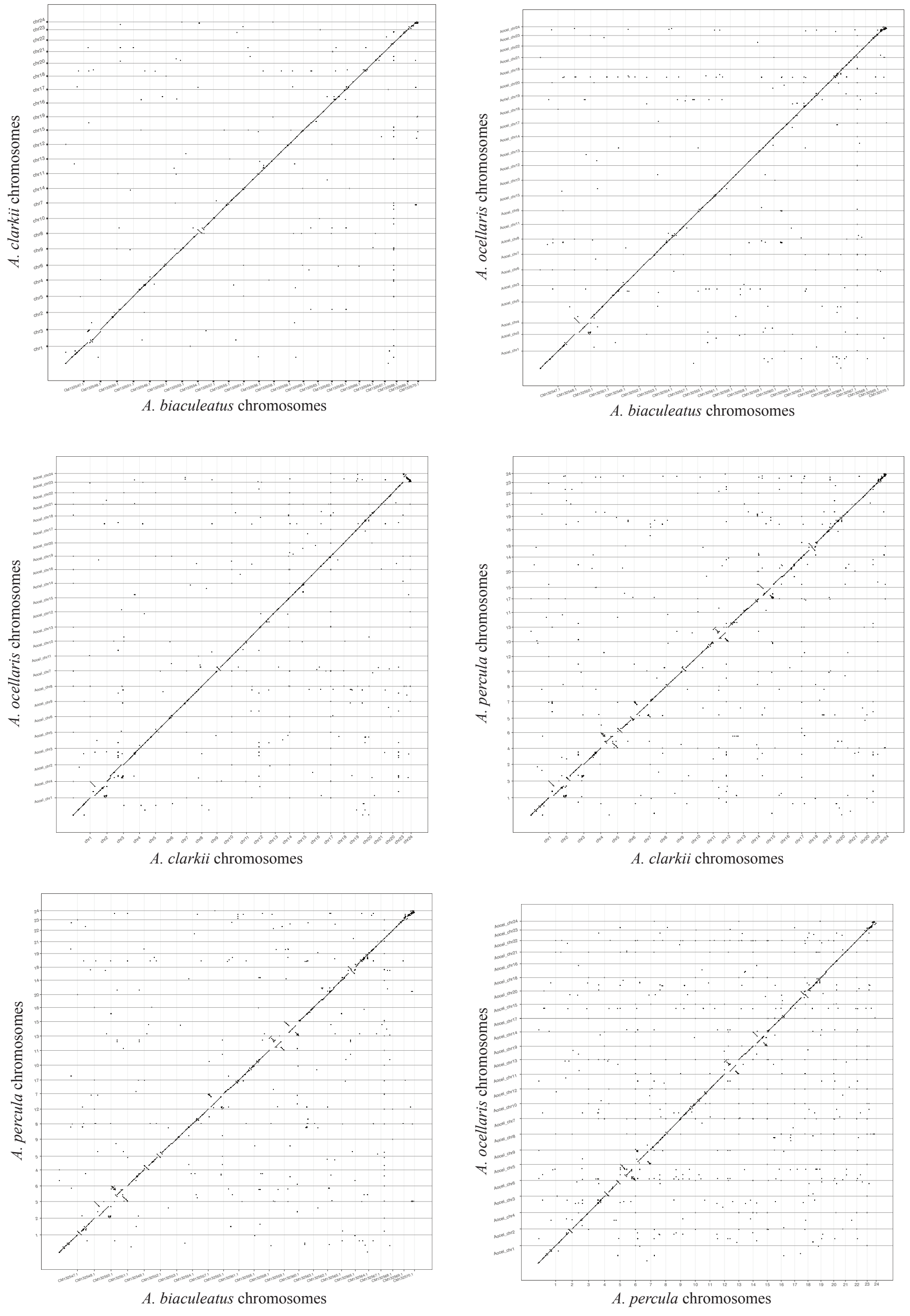


Figure S2: Whole-genome alignment dot plots between the four publicly available chromosome-level clownfish reference genomes: *A. biaculeatus* (GCA_053813585.1), *A. clarkii* (GCA_027123335.1), *A. ocellaris* (GCA_022539595.1), and *A. percula* (GCA_003047355.2). Each panel shows a pairwise comparison, with one genome on each axis. Alignments were performed with minimap2 (v2.28; `-x asm5`) and visualised with dotPlotly. The diagonal alignment pattern confirms a high degree of chromosomal synteny across all four genomes. Deviations from the diagonal indicate inversions or rearrangements between species. The published *A. percula* assembly shows extensive rearrangements relative to the other three genomes, consistent with its older sequencing data; an independent assembly was therefore generated for this species in the present study.

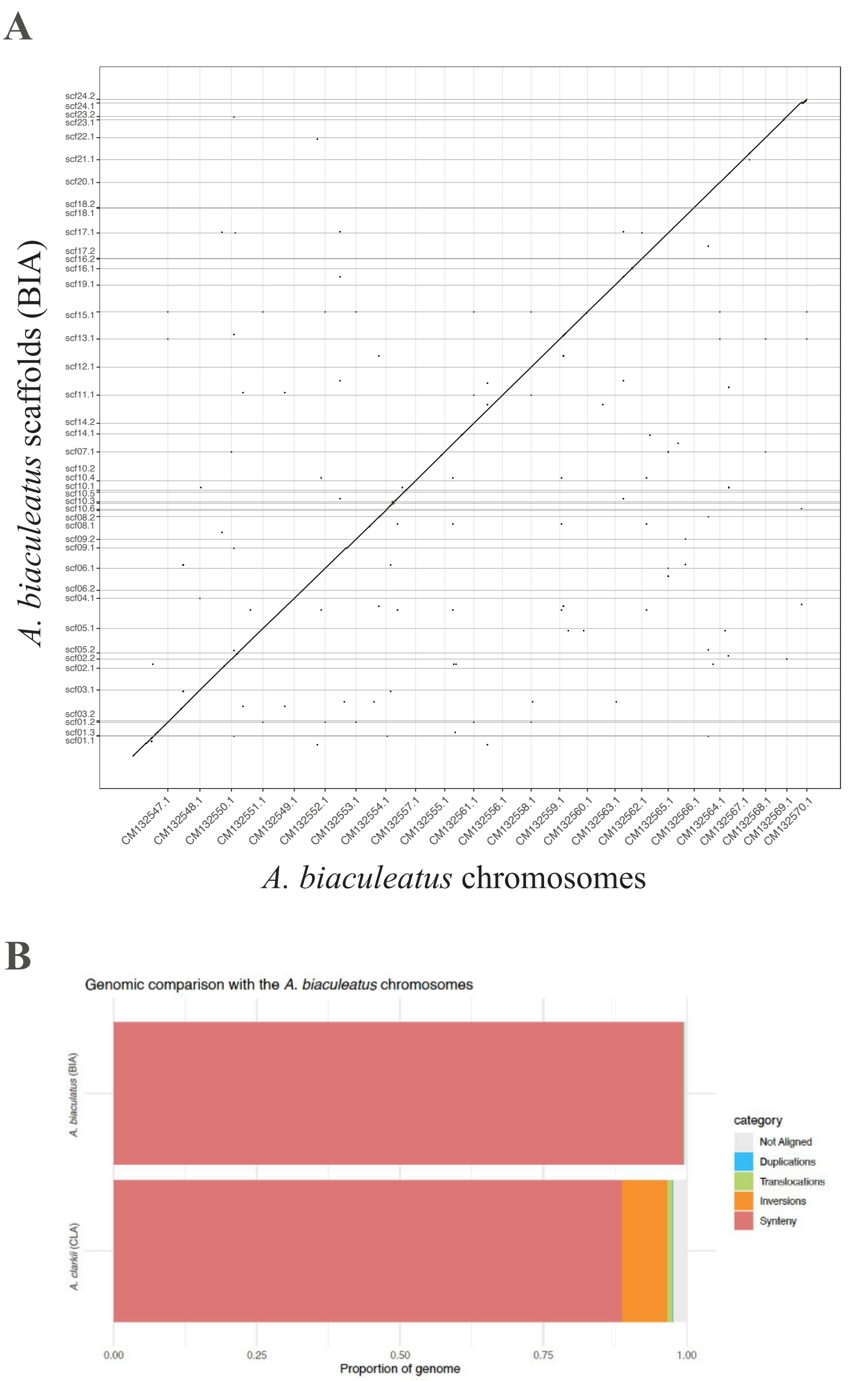


Figure S3: **A)** Whole-genome alignment dot plot between the *A. biaculeatus* assembly generated in this study (BIA; y-axis) and the publicly available chromosome-level reference genome (GCA_053813585.1; x-axis). Alignments were performed with minimap2 (v2.28; `-x asm5`) and visualised with dotPlotly. The near-perfect diagonal confirms that the two assemblies are highly concordant, with no major structural differences. **B)** Proportion of the genome classified as syntenic, not aligned, or affected by large structural variants (duplications, translocations, inversions) between the BIA assembly and the *A. biaculeatus* reference (top bar), as identified by SyRI. For comparison, the equivalent proportions for the pairwise comparison between *A. clarkii* (CLA) and the *A. biaculeatus* reference are also shown (bottom bar).

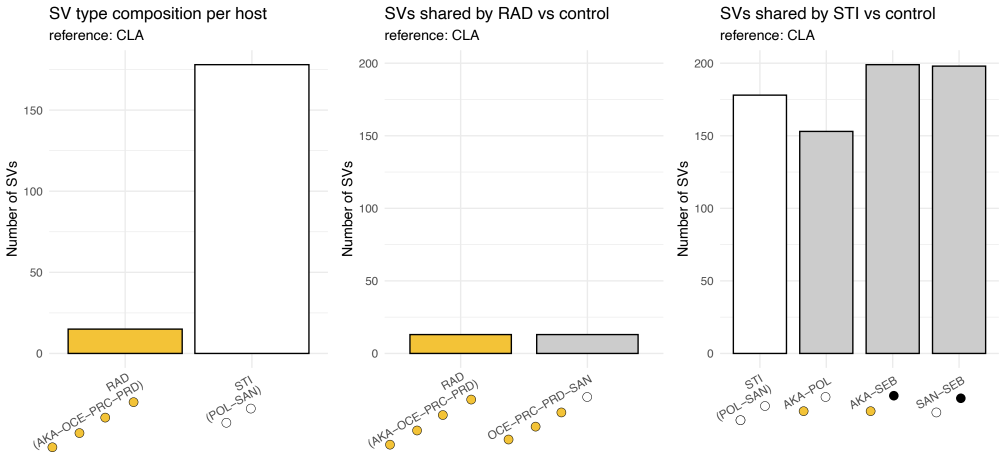


Figure S4: Host-specific structural variants (SVs) identified using *A. clarkii* (CLA) as the reference genome. Panels as in Supplementary Figure S28. **A)** SV counts per host category. **B)** RAD specialists versus control group. **C)** STI specialists versus control pairs. Results obtained using *A. clarkii* as reference are consistent with those obtained using *A. biaculeatus* (Supplementary Figure S28), indicating that the absence of a host-specialization signal is not an artefact of reference choice.

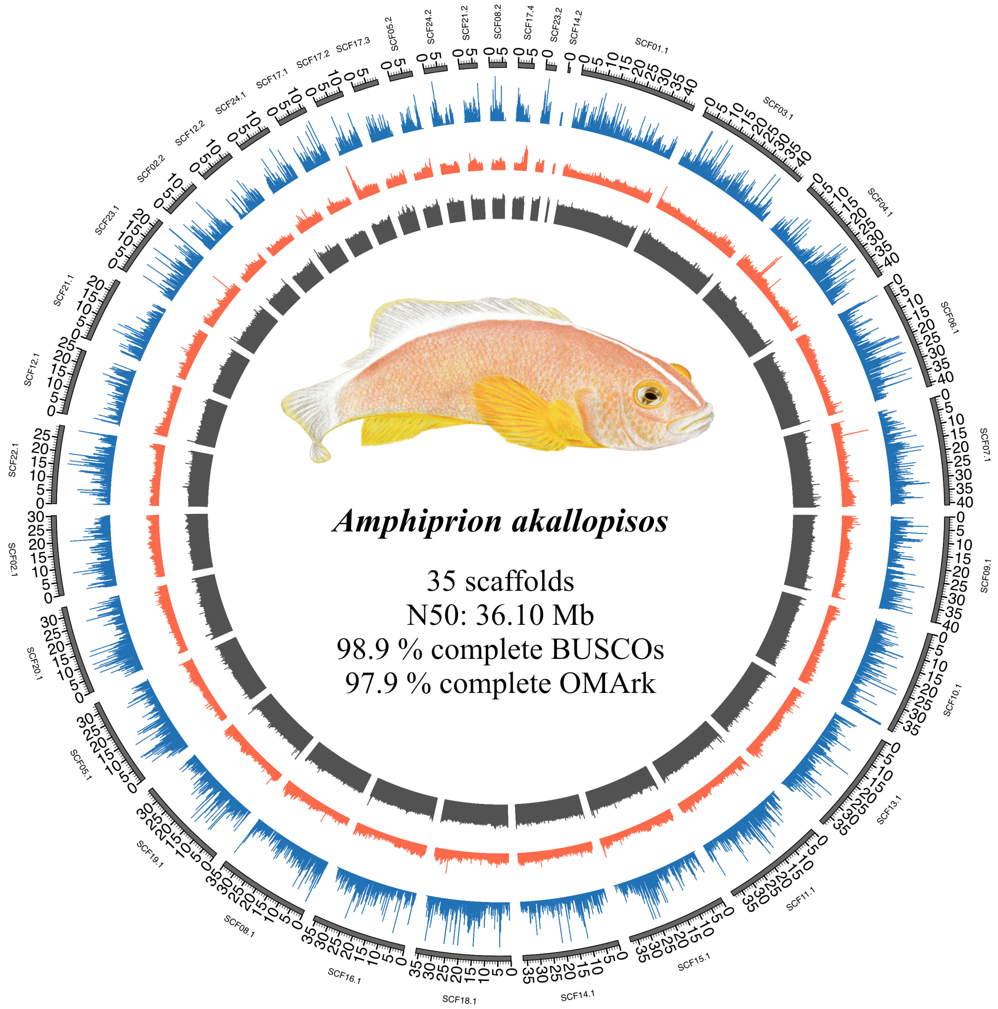


Figure S5: Genome assembly and annotation overview for *Amphiprion akallopisos*. The outer circle represents the assembled scaffolds. Tracks display gene density (blue), repeat density (red), and GC content (grey). Genome assembly completeness was assessed using BUSCO (v5.4.7; *actinopterygii_odb10* database), and genome annotation completeness was evaluated using OMArk (v0.3.1; Metazoa database). The circular plot was generated using Circos.

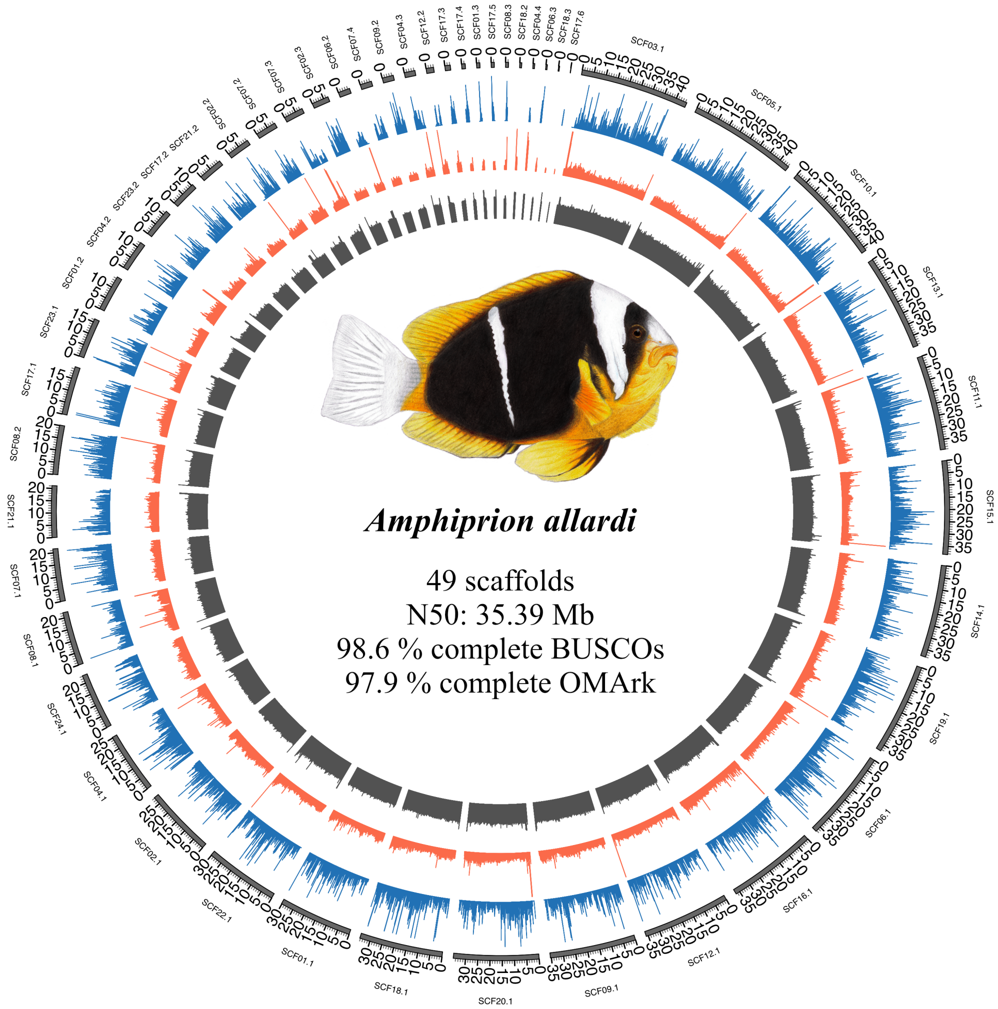


Figure S6: Genome assembly and annotation overview for *Amphiprion allardi*. The outer circle represents the assembled scaffolds. Tracks display gene density (blue), repeat density (red), and GC content (grey). Genome assembly completeness was assessed using BUSCO (v5.4.7; *actinopterygii_odb10* database), and genome annotation completeness was evaluated using OMArk (v0.3.1; Metazoa database). The circular plot was generated using Circos.

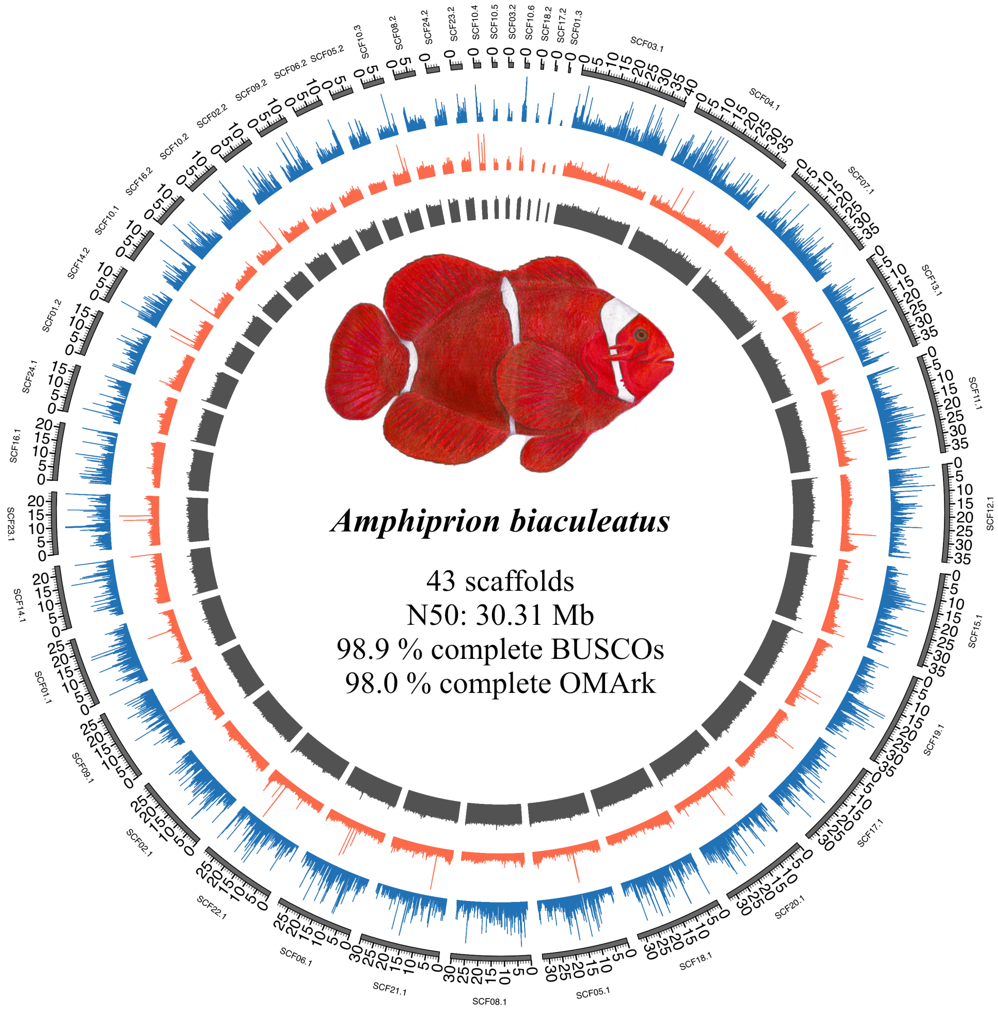


Figure S7: Genome assembly and annotation overview for *Amphiprion biaculeatus*. The outer circle represents the assembled scaffolds. Tracks display gene density (blue), repeat density (red), and GC content (grey). Genome assembly completeness was assessed using BUSCO (v5.4.7; *actinopterygii_odb10* database), and genome annotation completeness was evaluated using OMArk (v0.3.1; Metazoa database). The circular plot was generated using Circos.

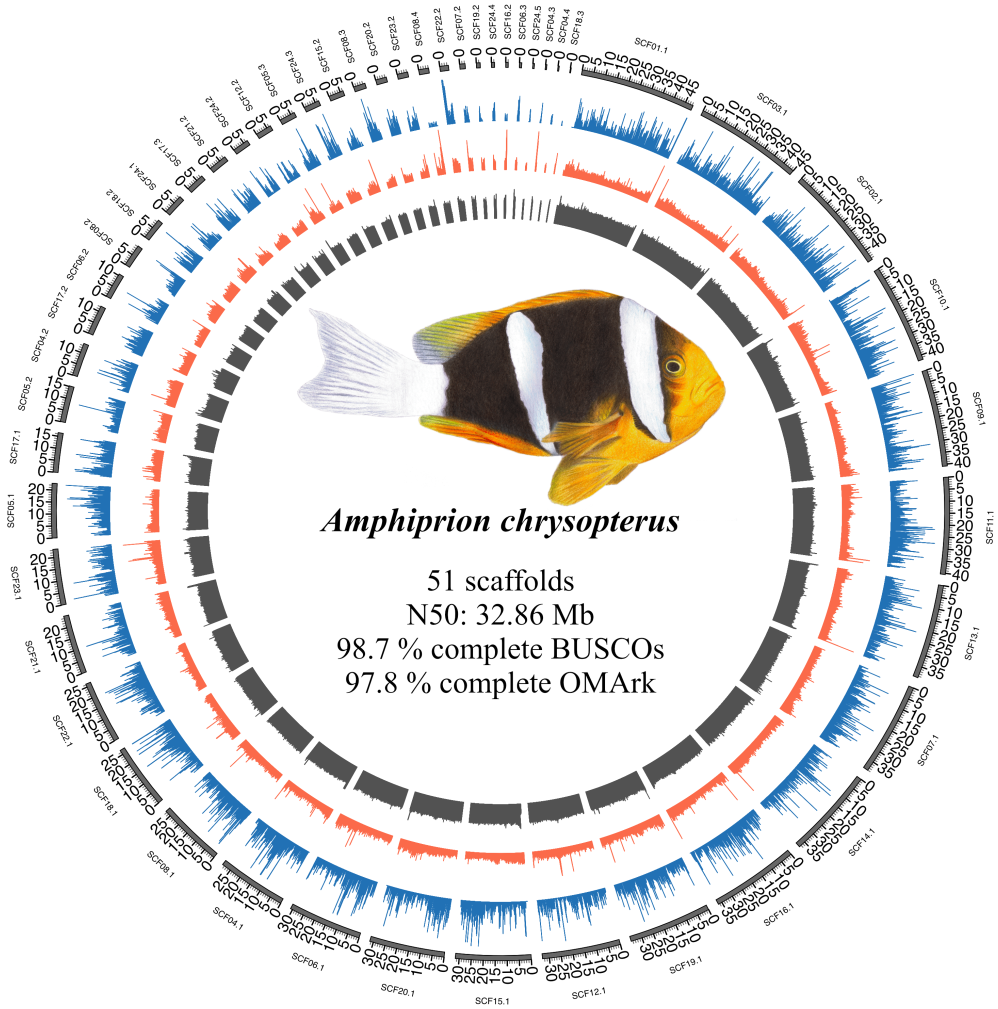


Figure S8: Genome assembly and annotation overview for *Amphiprion chrysopterus*. The outer circle represents the assembled scaffolds. Tracks display gene density (blue), repeat density (red), and GC content (grey). Genome assembly completeness was assessed using BUSCO (v5.4.7; *actinopterygii_odb10* database), and genome annotation completeness was evaluated using OMArk (v0.3.1; Metazoa database). The circular plot was generated using Circos.

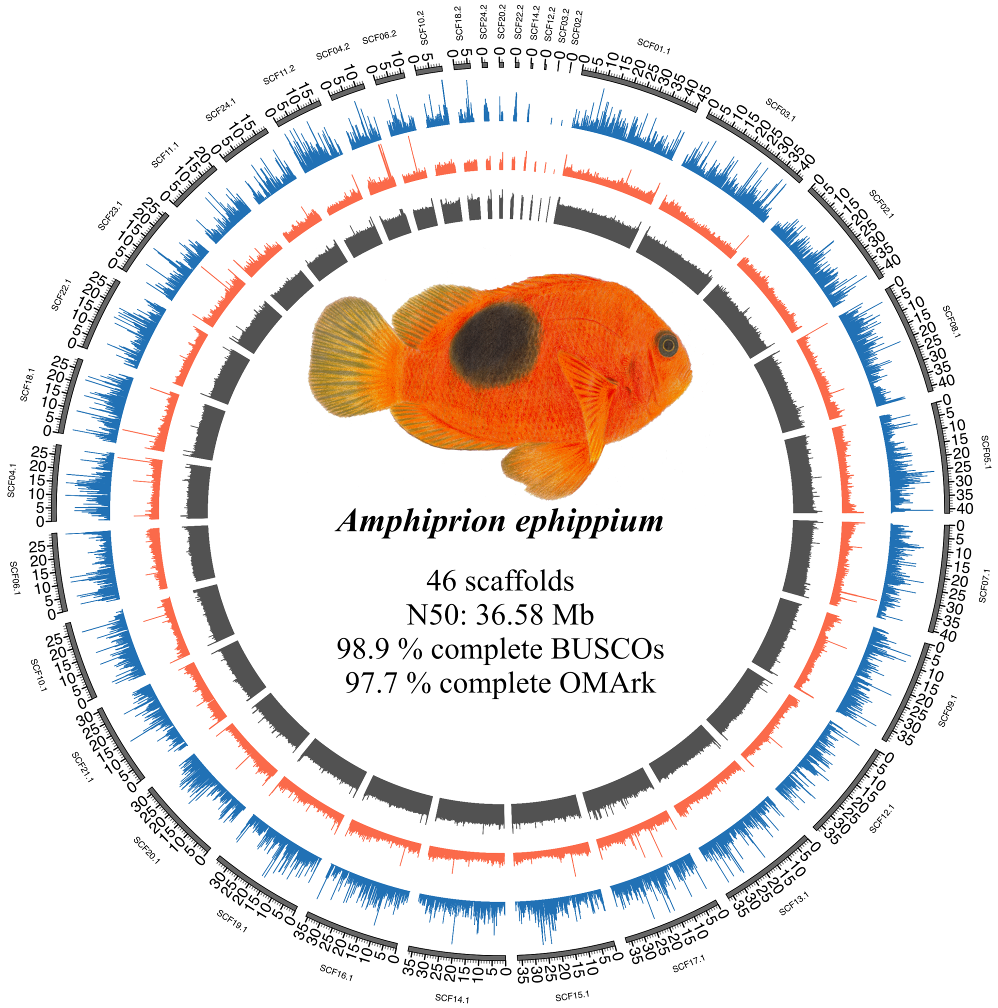


Figure S9: Genome assembly and annotation overview for *Amphiprion ephippium*. The outer circle represents the assembled scaffolds. Tracks display gene density (blue), repeat density (red), and GC content (grey). Genome assembly completeness was assessed using BUSCO (v5.4.7; *actinopterygii_odb10* database), and genome annotation completeness was evaluated using OMArk (v0.3.1; Metazoa database). The circular plot was generated using Circos.

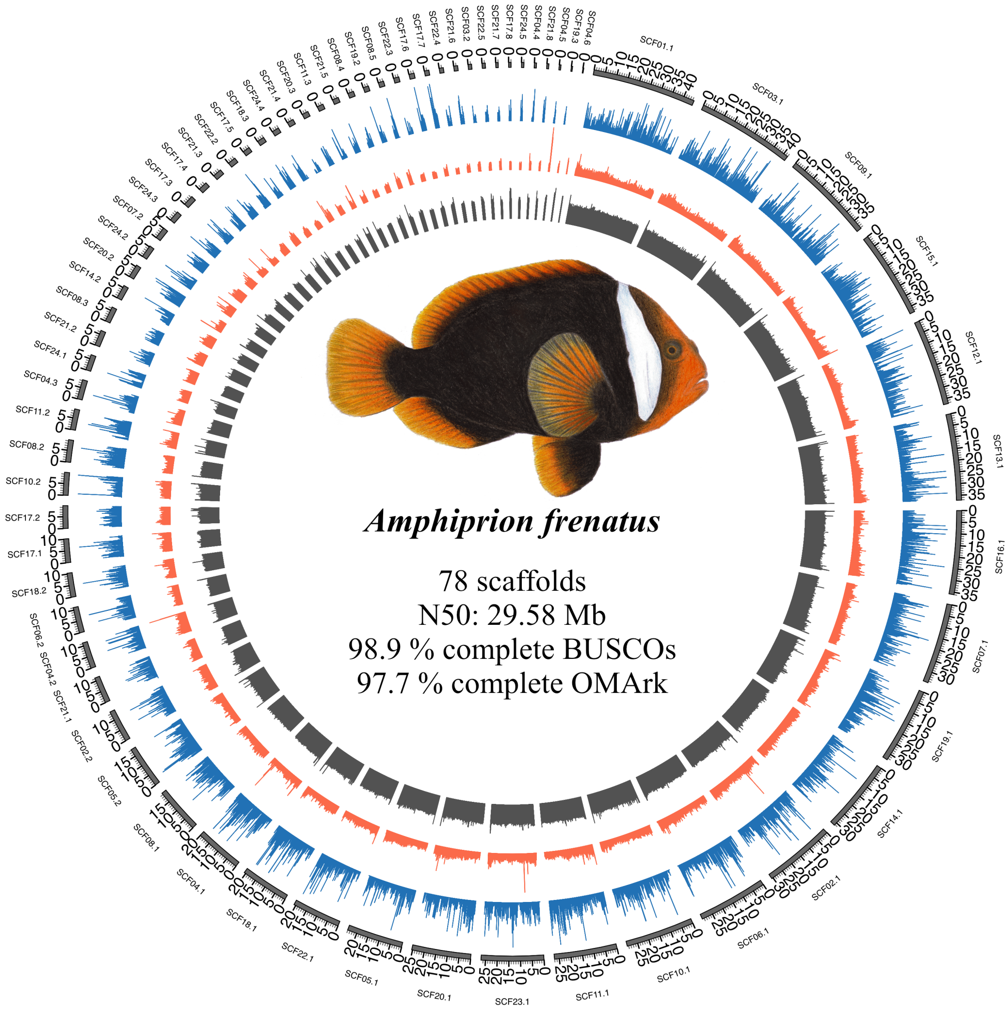


Figure S10: Genome assembly and annotation overview for *Amphiprion frenatus*. The outer circle represents the assembled scaffolds. Tracks display gene density (blue), repeat density (red), and GC content (grey). Genome assembly completeness was assessed using BUSCO (v5.4.7; *actinopterygii_odb10* database), and genome annotation completeness was evaluated using OMArk (v0.3.1; Metazoa database). The circular plot was generated using Circos.

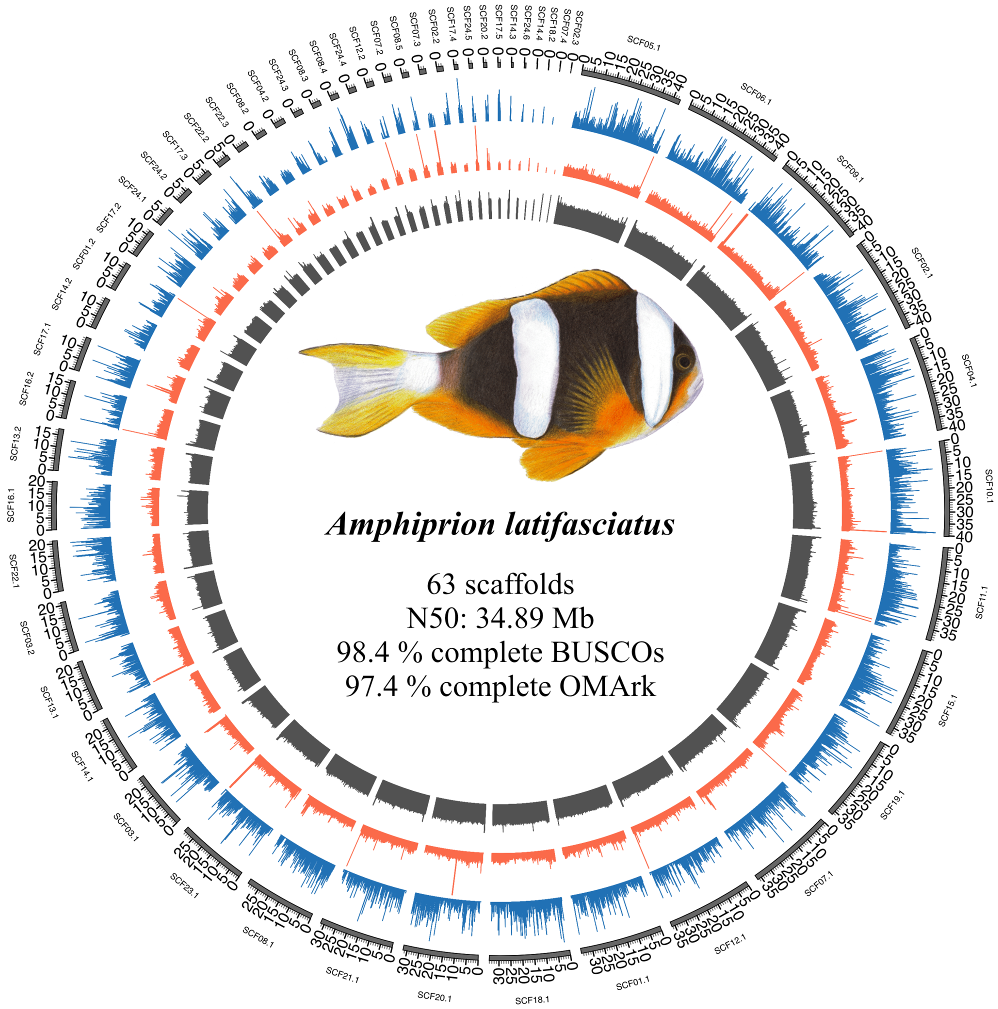


Figure S11: Genome assembly and annotation overview for *Amphiprion latifasciatus*. The outer circle represents the assembled scaffolds. Tracks display gene density (blue), repeat density (red), and GC content (grey). Genome assembly completeness was assessed using BUSCO (v5.4.7; *actinopterygii_odb10* database), and genome annotation completeness was evaluated using OMArk (v0.3.1; Metazoa database). The circular plot was generated using Circos.

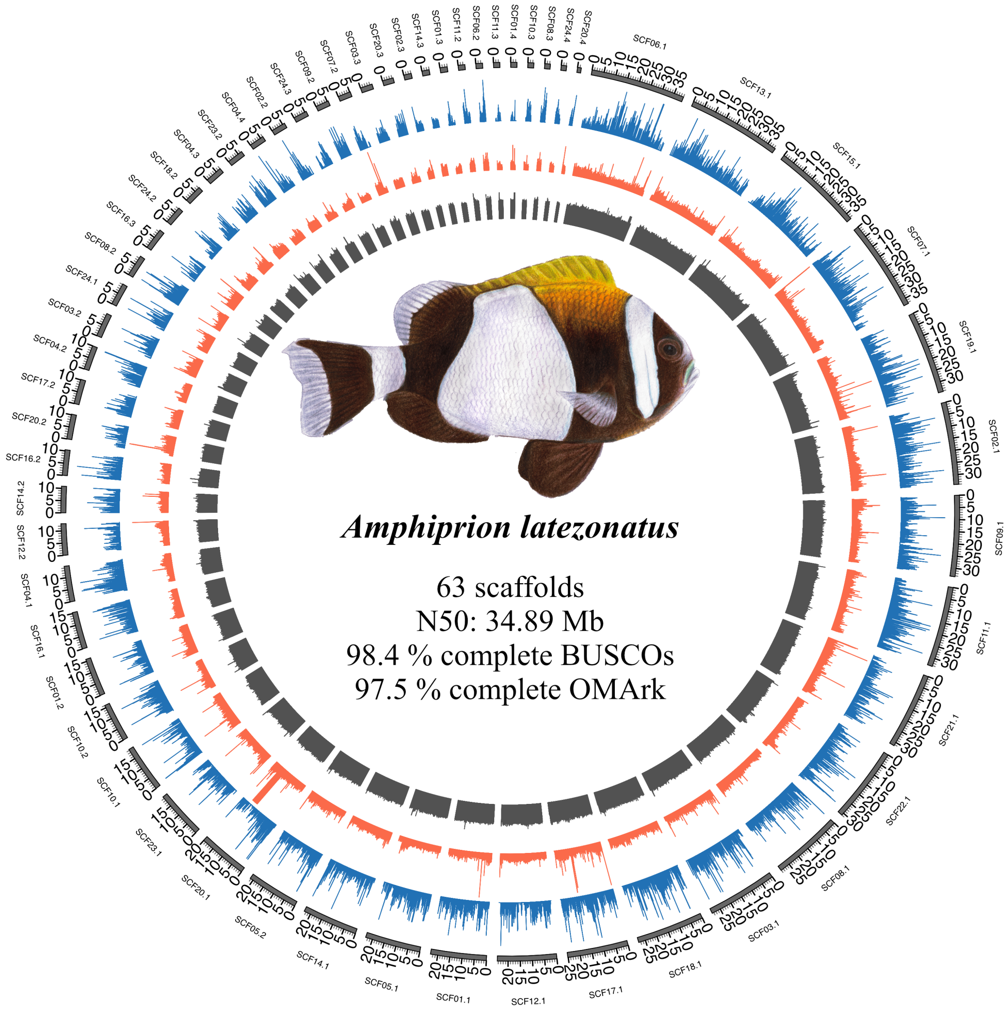


Figure S12: Genome assembly and annotation overview for *Amphiprion latezonatus*. The outer circle represents the assembled scaffolds. Tracks display gene density (blue), repeat density (red), and GC content (grey). Genome assembly completeness was assessed using BUSCO (v5.4.7; *actinopterygii_odb10* database), and genome annotation completeness was evaluated using OMArk (v0.3.1; Metazoa database). The circular plot was generated using Circos.

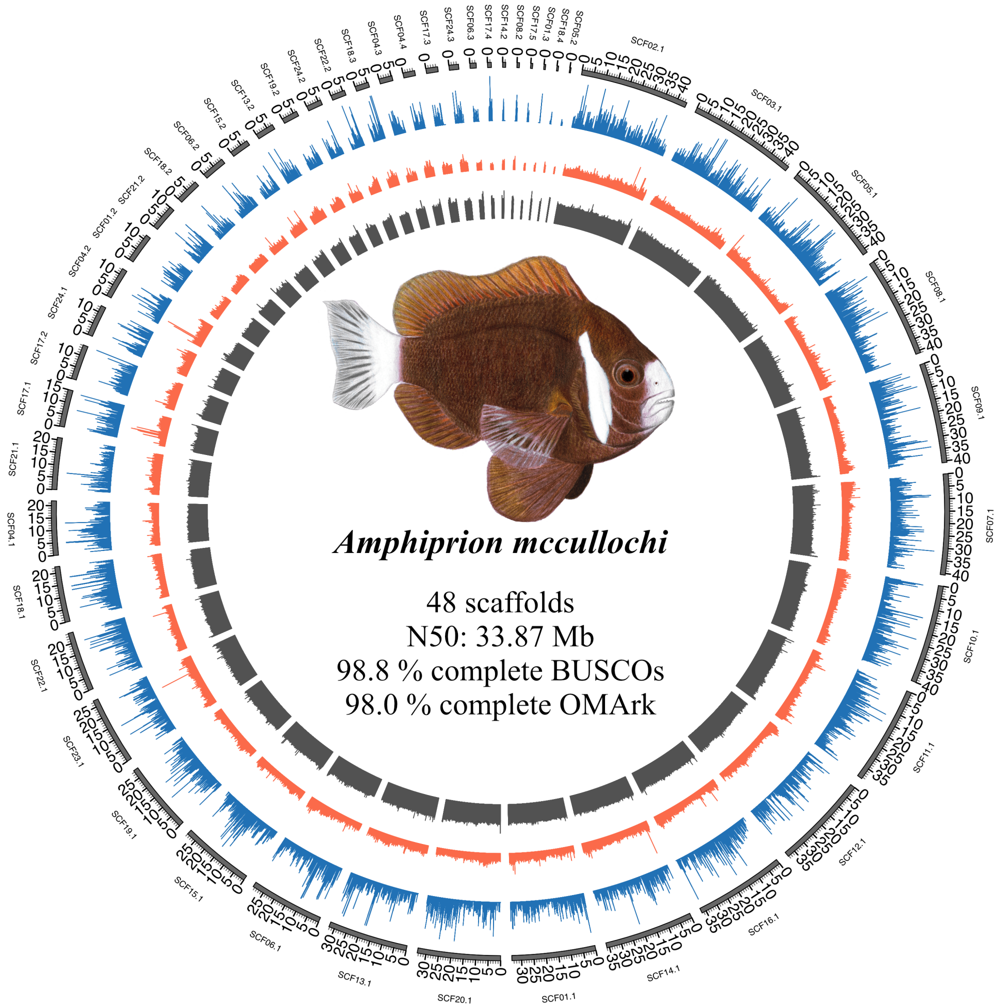


Figure S13: Genome assembly and annotation overview for *Amphiprion mccullochi*. The outer circle represents the assembled scaffolds. Tracks display gene density (blue), repeat density (red), and GC content (grey). Genome assembly completeness was assessed using BUSCO (v5.4.7; *actinopterygii_odb10* database), and genome annotation completeness was evaluated using OMArk (v0.3.1; Metazoa database). The circular plot was generated using Circos.

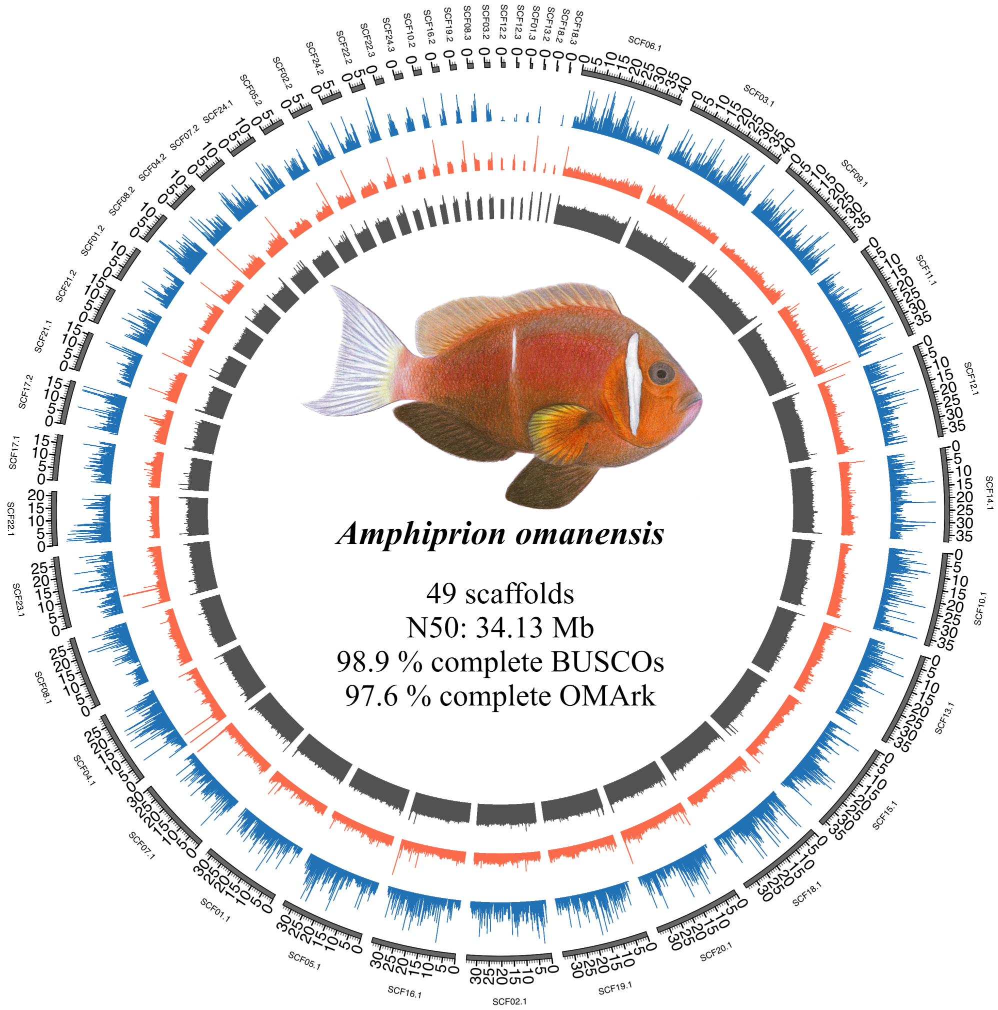


Figure S14: Genome assembly and annotation overview for *Amphiprion omanensis*. The outer circle represents the assembled scaffolds. Tracks display gene density (blue), repeat density (red), and GC content (grey). Genome assembly completeness was assessed using BUSCO (v5.4.7; *actinopterygii_odb10* database), and genome annotation completeness was evaluated using OMArk (v0.3.1; Metazoa database). The circular plot was generated using Circos.

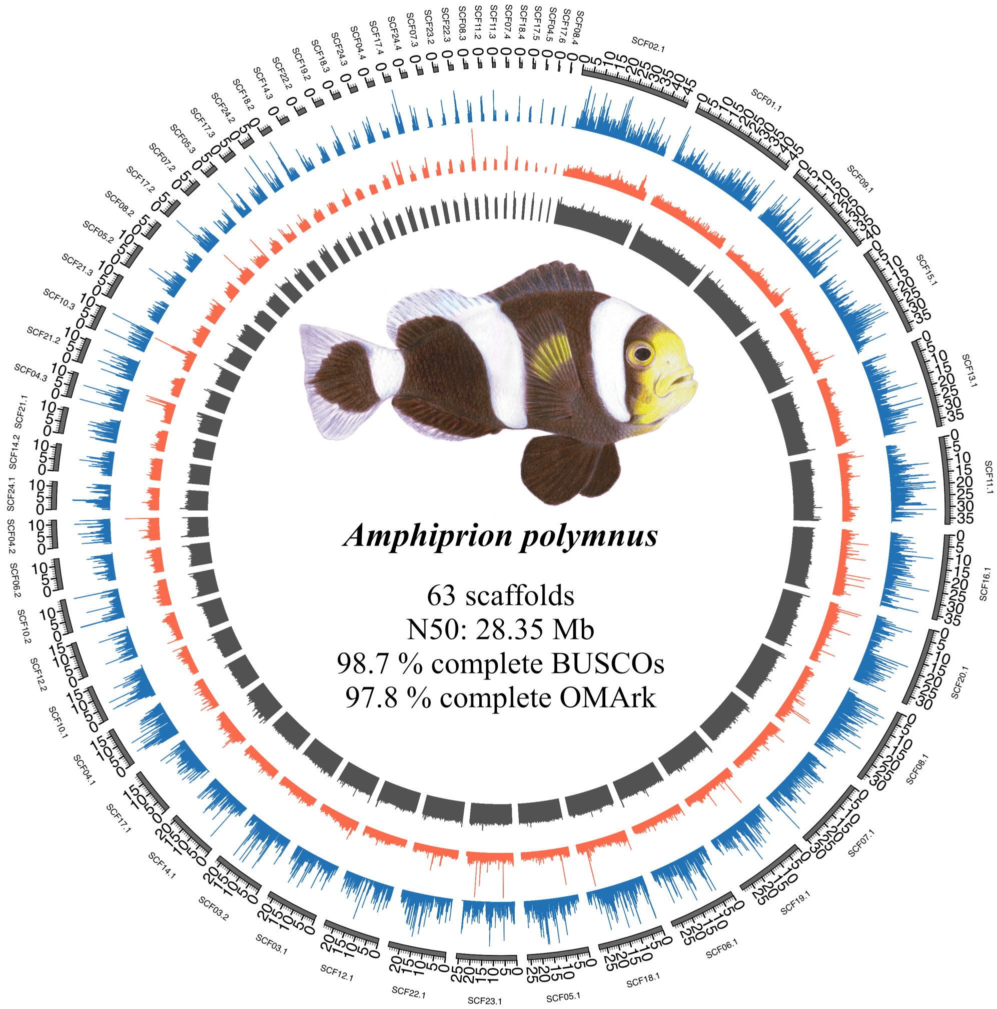


Figure S15: Genome assembly and annotation overview for *Amphiprion polymnus*. The outer circle represents the assembled scaffolds. Tracks display gene density (blue), repeat density (red), and GC content (grey). Genome assembly completeness was assessed using BUSCO (v5.4.7; *actinopterygii_odb10* database), and genome annotation completeness was evaluated using OMArk (v0.3.1; Metazoa database). The circular plot was generated using Circos.

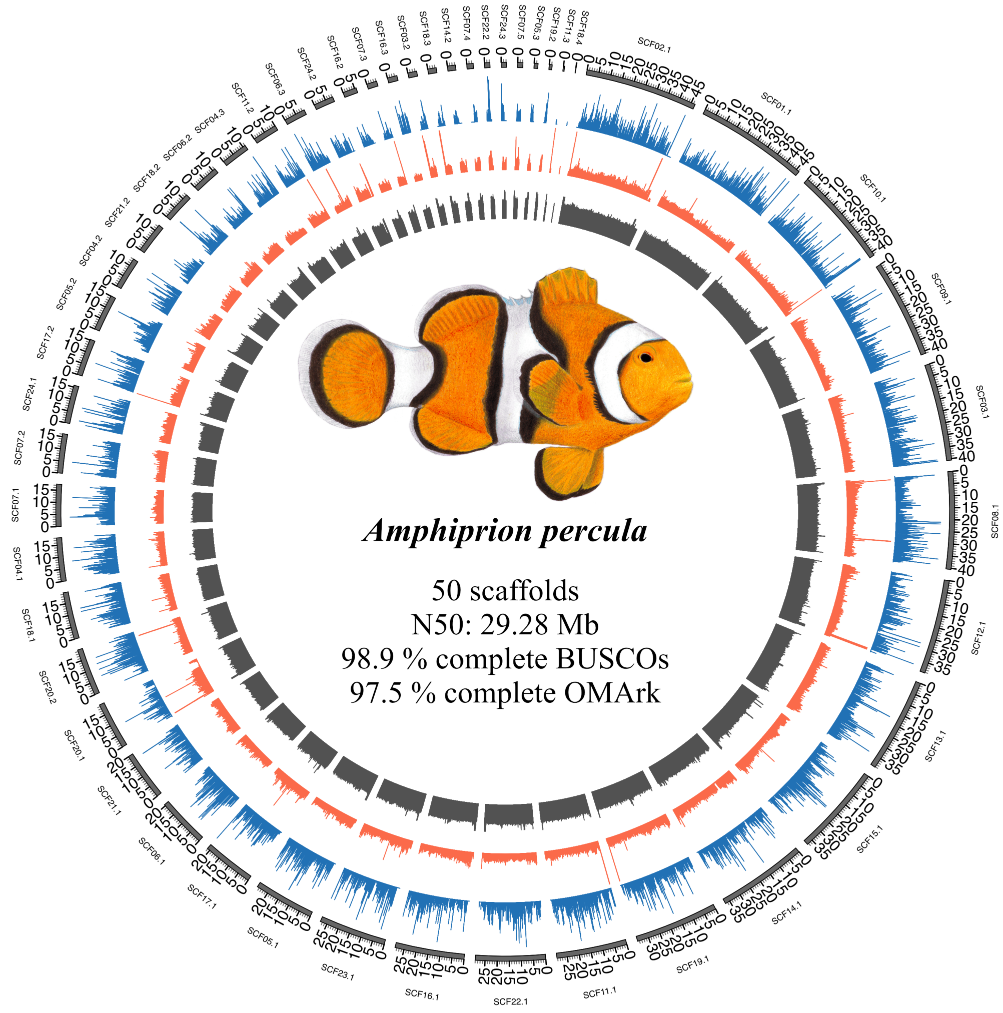


Figure S16: Genome assembly and annotation overview for *Amphiprion percula*. The outer circle represents the assembled scaffolds. Tracks display gene density (blue), repeat density (red), and GC content (grey). Genome assembly completeness was assessed using BUSCO (v5.4.7; *actinopterygii_odb10* database), and genome annotation completeness was evaluated using OMArk (v0.3.1; Metazoa database). The circular plot was generated using Circos.

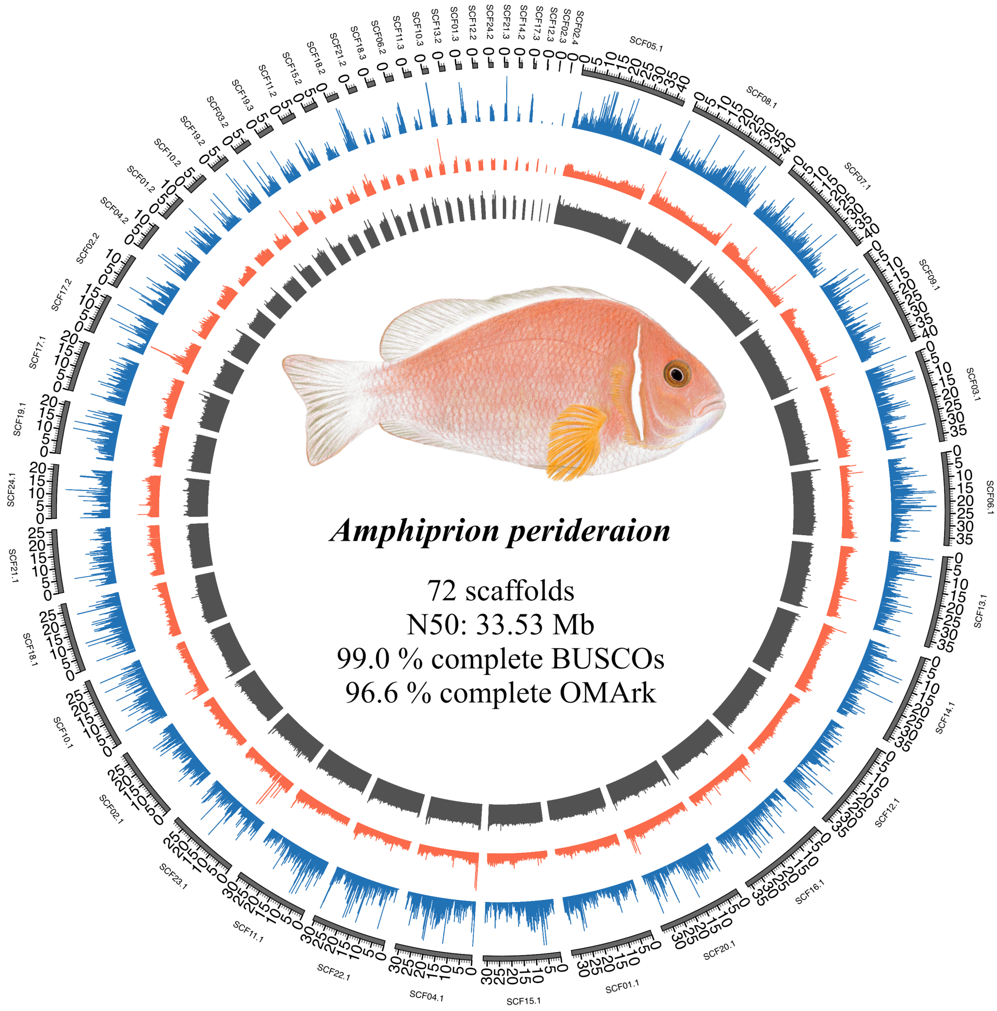


Figure S17: Genome assembly and annotation overview for *Amphiprion perideraion*. The outer circle represents the assembled scaffolds. Tracks display gene density (blue), repeat density (red), and GC content (grey). Genome assembly completeness was assessed using BUSCO (v5.4.7; *actinopterygii_odb10* database), and genome annotation completeness was evaluated using OMArk (v0.3.1; Metazoa database). The circular plot was generated using Circos.

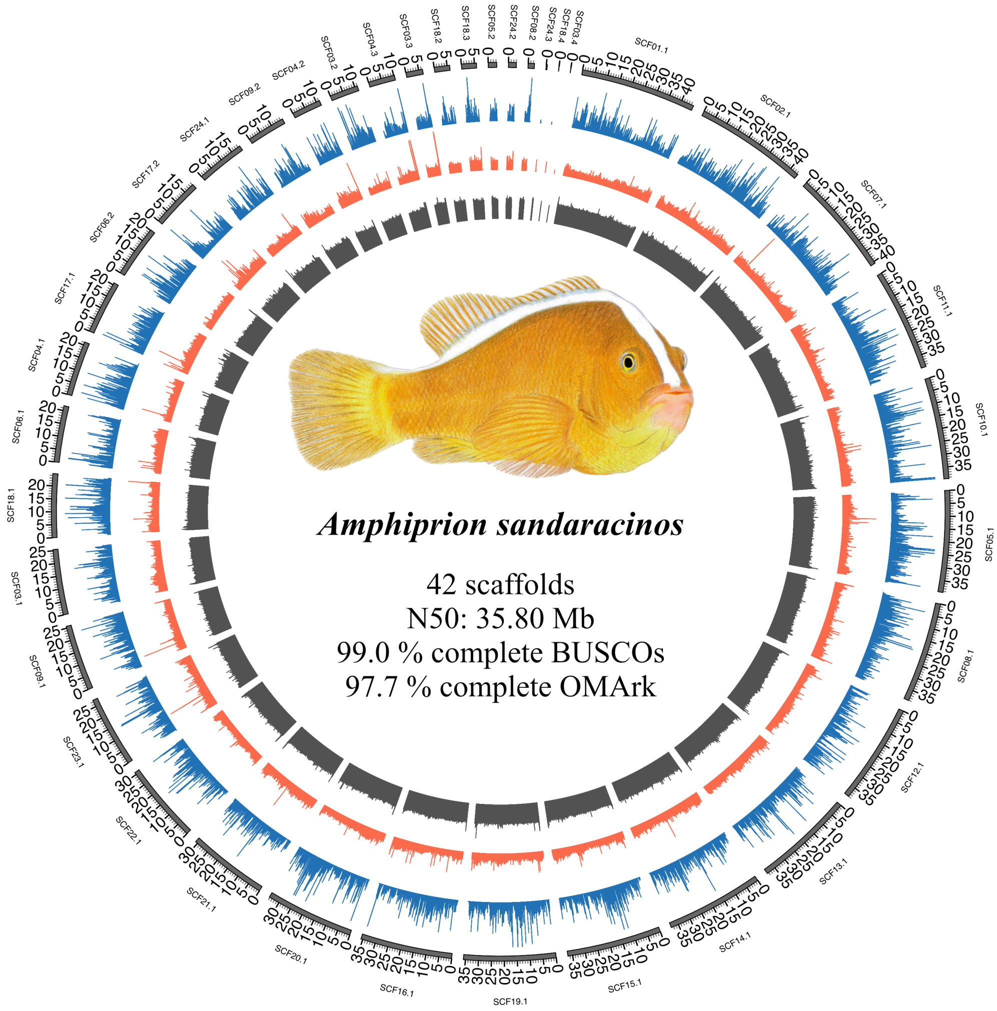


Figure S18: Genome assembly and annotation overview for *Amphiprion sandaracinos*. The outer circle represents the assembled scaffolds. Tracks display gene density (blue), repeat density (red), and GC content (grey). Genome assembly completeness was assessed using BUSCO (v5.4.7; *actinopterygii_odb10* database), and genome annotation completeness was evaluated using OMArk (v0.3.1; Metazoa database). The circular plot was generated using Circos.

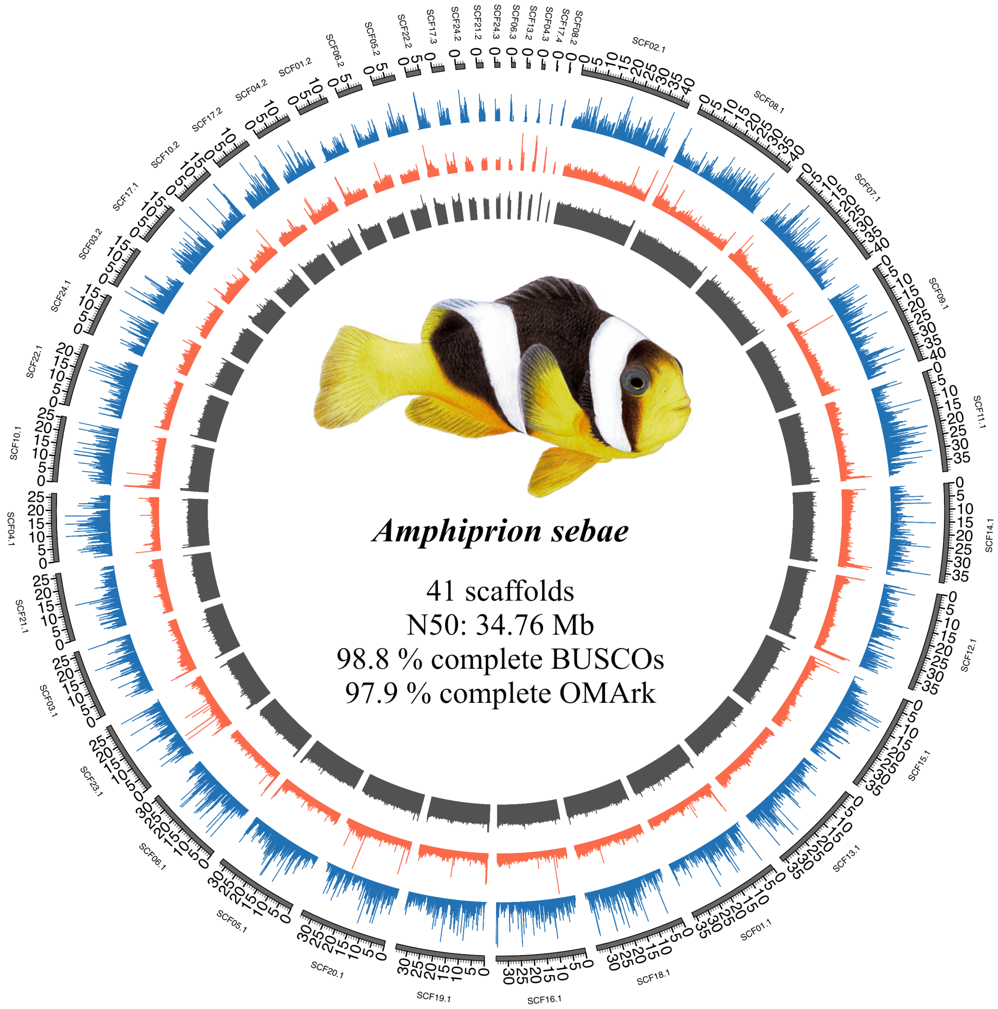


Figure S19: Genome assembly and annotation overview for *Amphiprion sebae*. The outer circle represents the assembled scaffolds. Tracks display gene density (blue), repeat density (red), and GC content (grey). Genome assembly completeness was assessed using BUSCO (v5.4.7; *actinopterygii_odb10* database), and genome annotation completeness was evaluated using OMArk (v0.3.1; Metazoa database). The circular plot was generated using Circos.

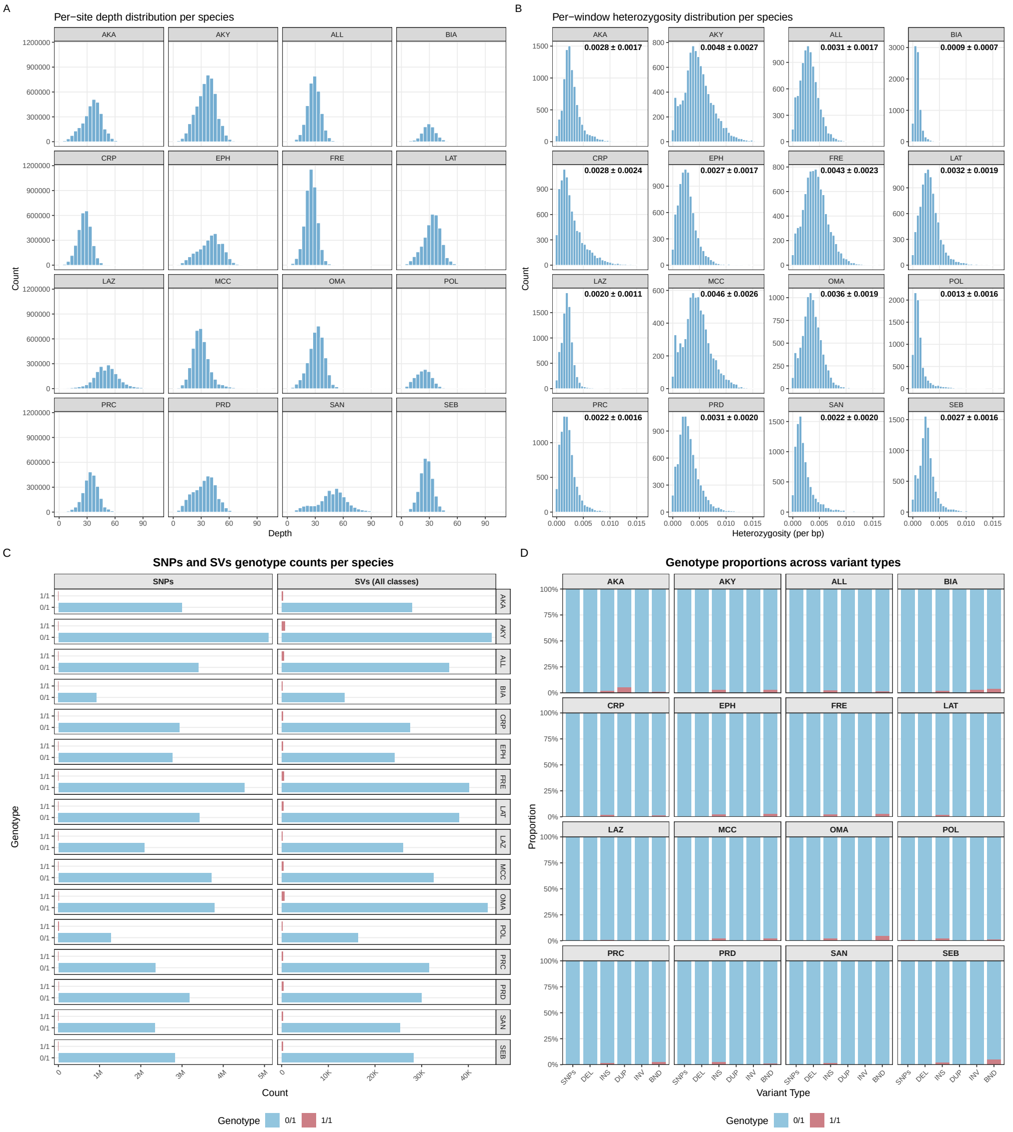


Figure S20: Depth and heterozygosity based on reads mapping to the corresponding assembly. **A)** Per-site sequencing depth distribution for each species, based on HiFi reads mapped back to their corresponding assembly. **B)** Per-window heterozygosity distribution for each species, with mean $\pm$ standard deviation indicated above each panel. **C)** Genotype counts for SNPs (left) and SVs of all classes combined (right) for each species, distinguishing heterozygous (0/1) from homozygous-alternate (1/1) genotypes. **D)** Proportion of heterozygous (0/1) versus homozygous-alternate (1/1) genotypes for SNPs and for each SV class (DEL, INS, DUP, INV, BND) in each species. Across all species, the large majority of called variants were heterozygous, and homozygous-alternate genotypes were rare, consistent with high accuracy and low error rate of the assemblies.

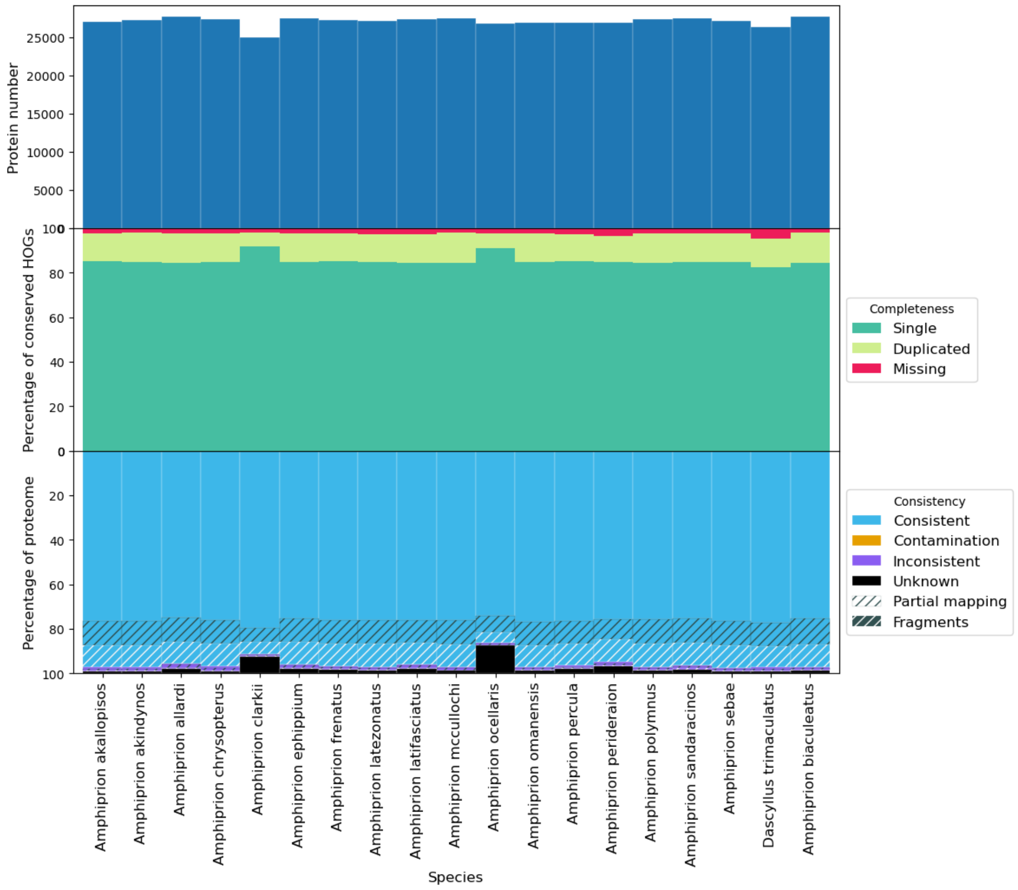


Figure S21: Annotation completeness assessed with OMArk (v0.3.1; Metazoa database) for all 16 newly assembled species, together with the publicly available assemblies of *A. ocellaris* and *A. clarkii* included for comparison. The upper panel shows the percentage of conserved Hierarchical Orthologous Groups (HOGs) detected as single-copy, duplicated, or missing for each species. The lower panel shows the consistency of the predicted proteome: the proportion of proteins classified as consistent (blue), unknown (black), inconsistent (purple), contamination orange), partially mapping (hatched), or fragments (hatched dark) relative to the Metazoa reference.

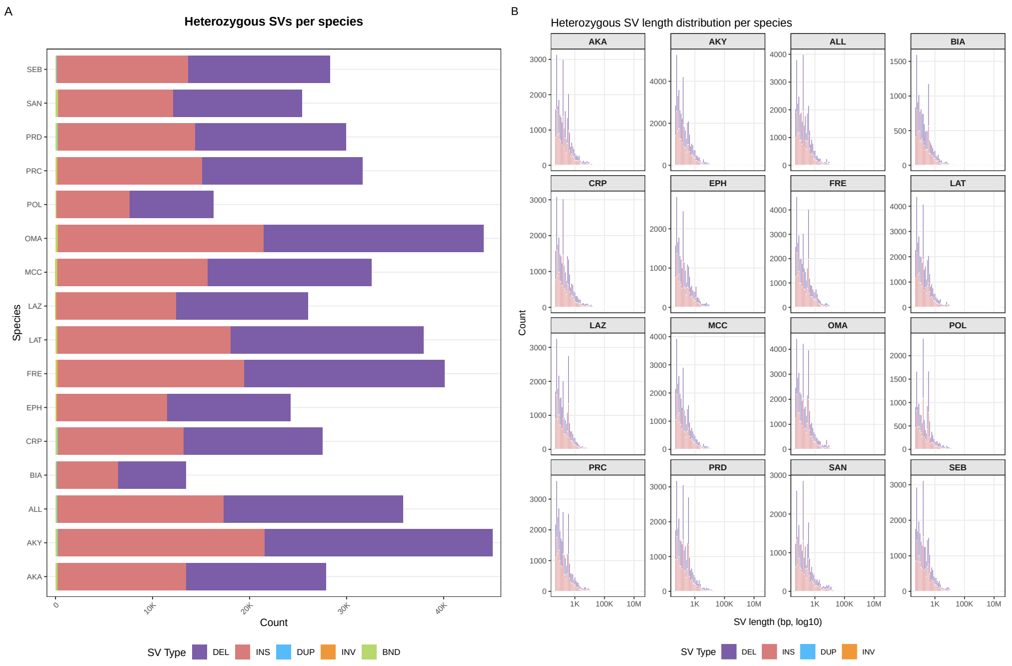


Figure S22: Heterozygous structural variants (SVs) identified within each individual assembly. textbfA) Number of heterozygous SVs per species, broken down by SV class (DEL, INS, DUP, INV, BND). **B)** Length distribution (bp, $\log_{10}$ scale) of heterozygous SVs per species, broken down by SV class. Across all species, heterozygous SVs were predominantly short insertions and deletions.

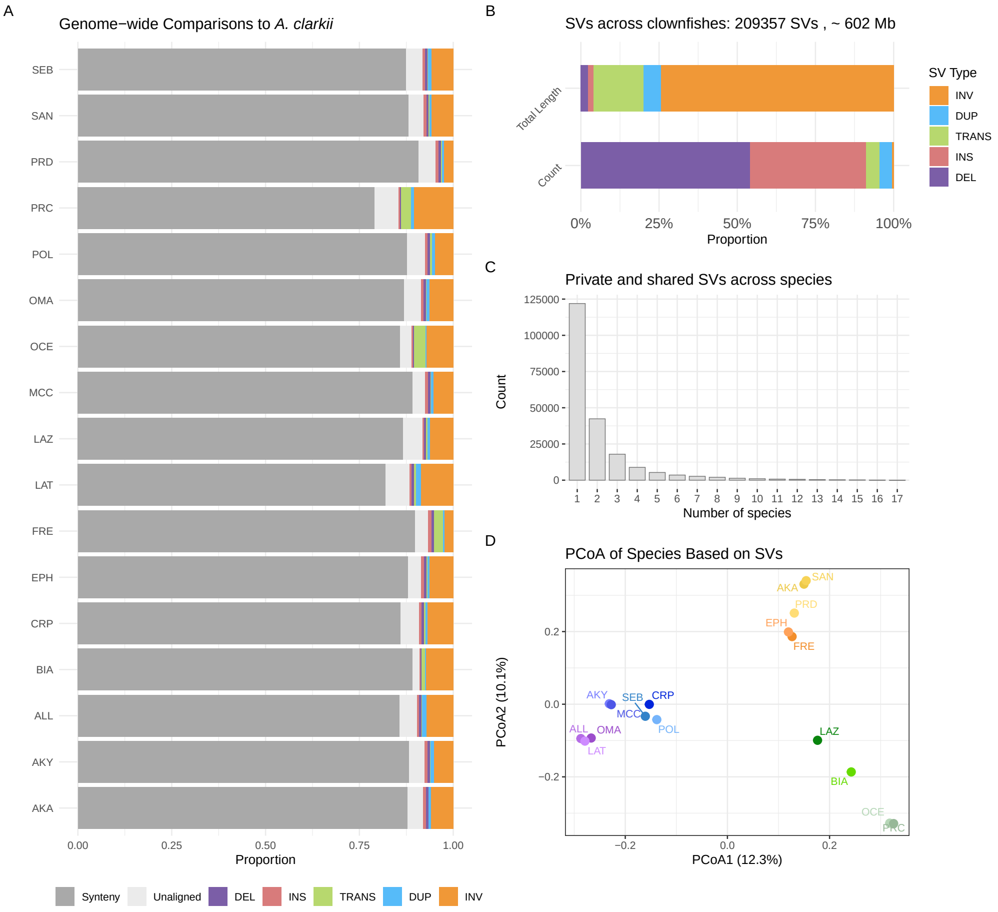


Figure S23: Structural variation across clownfish genomes relative to *A. clarkii*, analogous to Fig. 2.**A)** Proportion of each genome classified as syntenic, not aligned, or affected by each SV class (deletions, DEL; insertions, INS; translocations, TRANS; duplications, DUP; inversions, INV), relative to *A. clarkii*. **B)** Total count (bottom) and cumulative sequence length (top) of each SV class after merging SVs across species. A total of 209,357 SVs were identified, corresponding to a cumulative length of approximately 602 Mb; as with the *A. biaculeatus*-referenced dataset (Fig. 2B), INDELs are the most frequent class by count, while inversions dominate by cumulative length. **C)** Number of SVs as a function of the number of species sharing them. As with the *A. biaculeatus*-referenced dataset, most SVs are private, and the number of shared SVs decreases with the number of species sharing them. **D)** Principal coordinates analysis (PCoA) based on pairwise Jaccard distances computed from SV presence/absence across species (PCoA1: 12.3%, PCoA2: 10.1% of variance explained). As with the *A. biaculeatus*-referenced results (Fig. 2D), species cluster primarily by phylogenetic relationships rather than sea anemone host specialization.

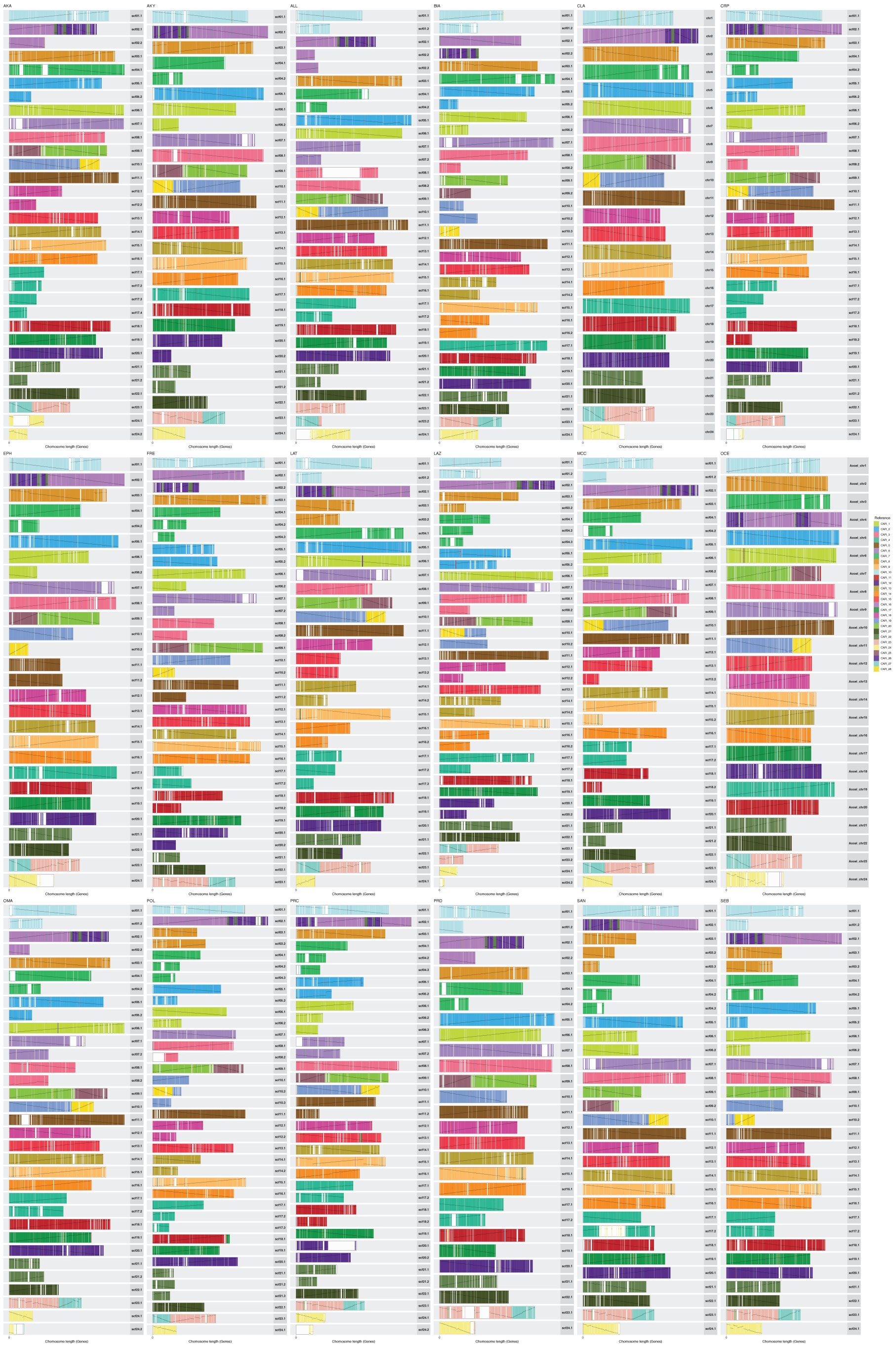


Figure S24: Synteny between extant clownfish genomes and the reconstructed clownfish ancestral scaffolds. For each species, scaffolds are represented as horizontal bars and coloured according to the homologous ancestral scaffold as inferred from Hierarchical Orthologous Groups (HOGs) using AGORA. Colours correspond to the 24 ancestral chromosomal units (reference colour key shown in the OCE panel). Inverted orientation of syntenic blocks relative to the ancestor is indicated. Only scaffolds or chromosomes containing more than 200 genes were considered. The publicly available *A. ocellaris* (OCE) and *A. clarkii* (CLA) chromosome-level assemblies are included for reference. Overall, synteny is highly conserved across species, with most ancestral chromosomal blocks recovered intact despite some fragmentation in the assemblies.

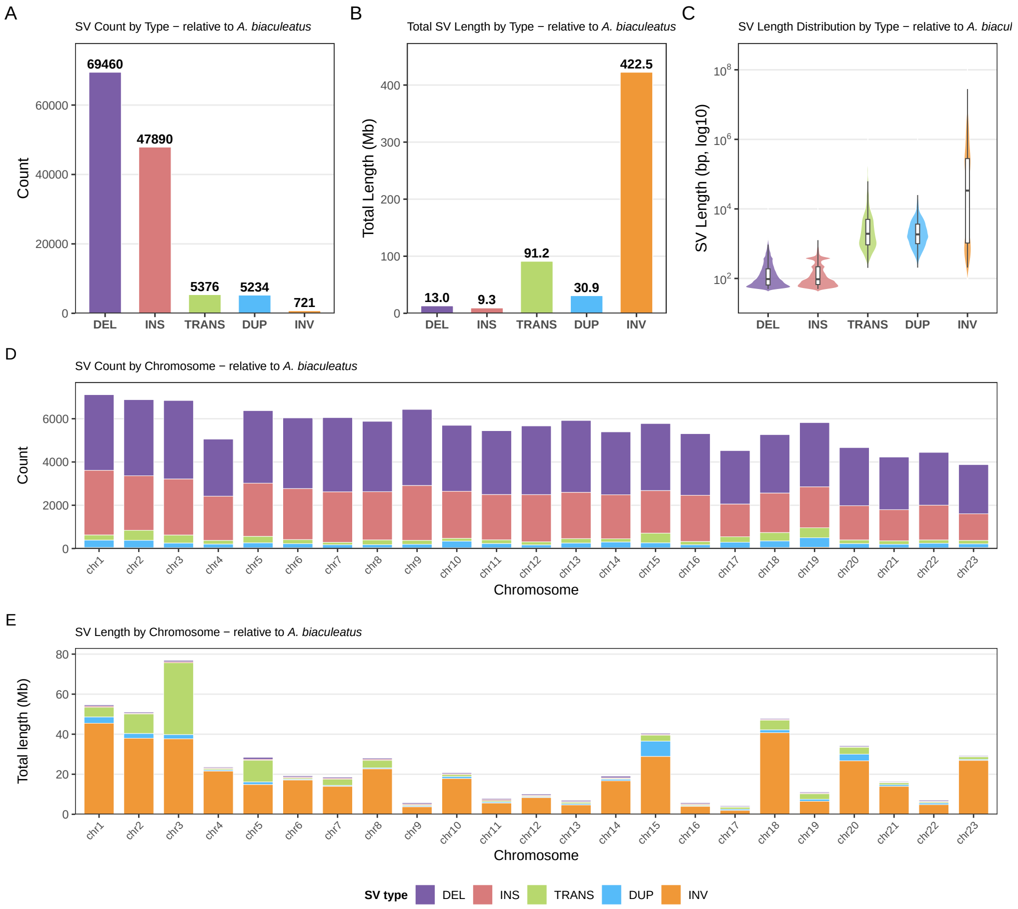


Figure S25: Detailed structural variant (SV) statistics for all species relative to *A. biaculeatus*. **A)** Total count of each SV class (deletions, DEL; insertions, INS; translocations, TRANS; duplications, DUP; inversions, INV) summed across all species after merging (see Fig. 2). **B)** Cumulative sequence length (Mb) affected by each SV class. **C)** Distribution of SV lengths (bp, $\log_{10}$ scale) for each class, showing that INDELs are typically short whereas inversions are considerably longer. **D)** Number of SVs per chromosome and **E)** cumulative sequence length of SVs per chromosome, broken down by SV class. SVs were detected in all chromosomes

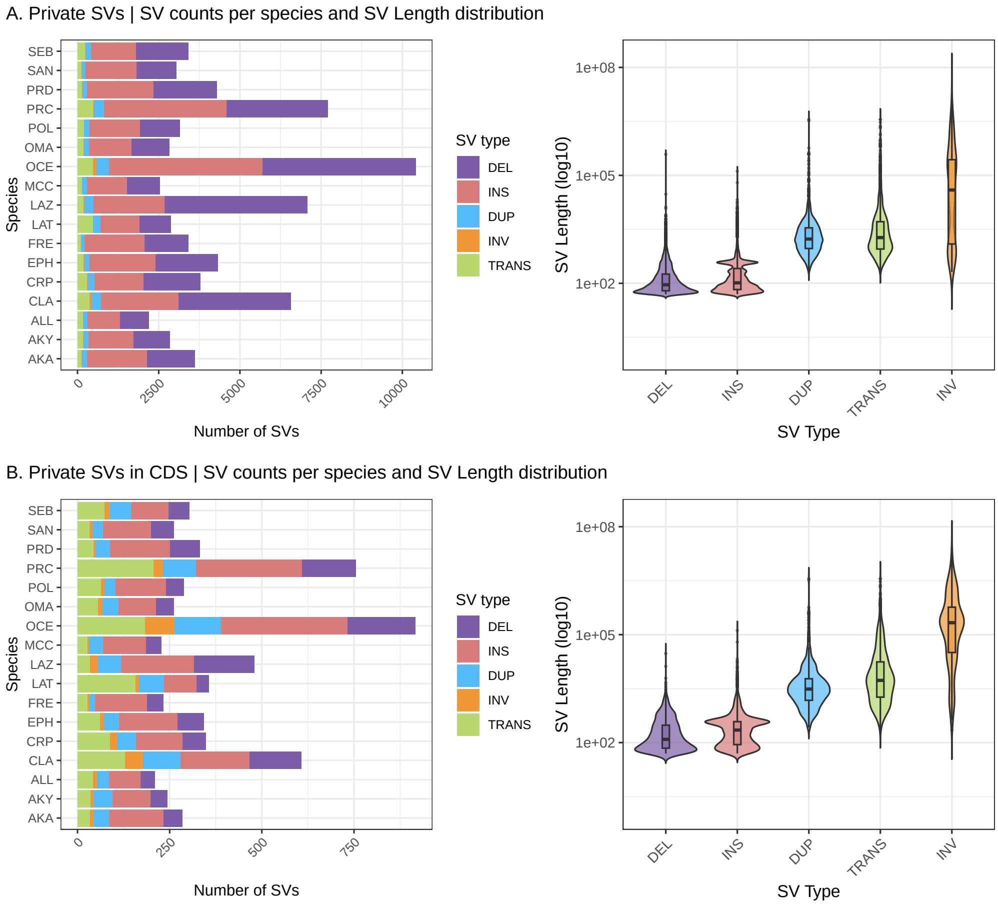


Figure S26: Private structural variants (SVs; i.e. detected in a single species only) relative to *A. biaculeatus*. **A)** Left: number of private SVs per species, broken down by SV class (DEL, INS, DUP, INV, TRANS). Right: length distribution (bp, $\log_{10}$ scale) of private SVs by class; as with the full SV catalogue (Fig. 2B), duplications, translocations and especially inversions were considerably longer on average than deletions and insertions. **B)** As in (A), restricted to private SVs overlapping protein-coding sequences (CDS). Private CDS-overlapping SVs were far less numerous than the full private SV set.

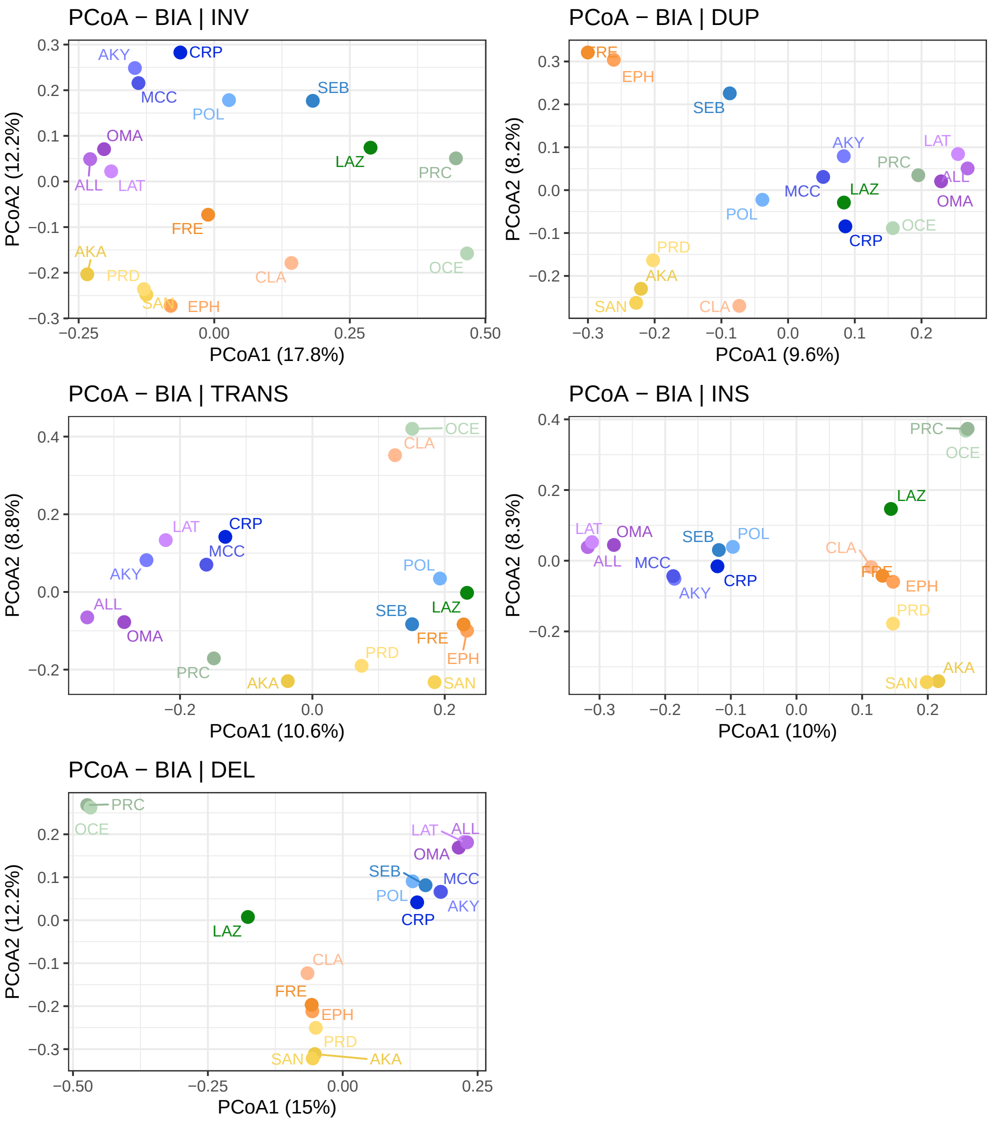


Figure S27: Principal coordinates analysis (PCoA) of species based on the presence/absence of each structural variant (SV) class separately: inversions (INV), duplications (DUP), translocations (TRANS), insertions (INS), and deletions (DEL). For each class, pairwise Jaccard distances were computed across species and visualised using `cmdscale`. The percentage of variance explained by PCoA1 and PCoA2 is indicated on each axis. In all cases, species cluster primarily according to their phylogenetic relationships, with no clear grouping driven by sea anemone host specialization detectable on the first axes of variation. Species abbreviations follow Fig. 2.

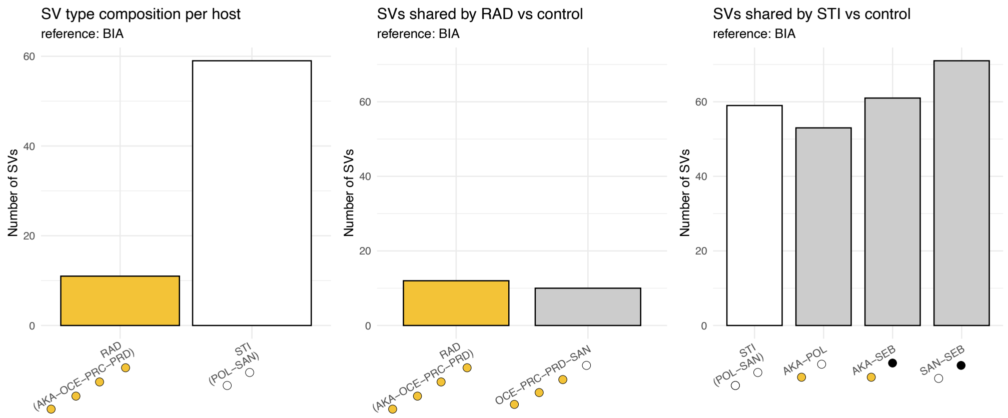


Figure S28: Host-specific structural variants (SVs) identified using *A. biaculeatus* (BIA) as the reference genome. **A)** Number of SVs shared among all members of two host-specialist groups: *Radianthus* specialists (RAD; *A. akallopisos*, *A. ocellaris*, *A. percula*, and *A. perideraion*) and *Stichodactyla* specialists (STI; *A. polymnus* and *A. sandaracinos*). **B)** Number of SVs shared among RAD specialists compared with an equally sized control group of species with different host associations (*A. ocellaris*, *A. percula*, *A. perideraion*, and *A. sandaracinos*). **C)** Number of SVs shared between STI specialists compared with three control species pairs that do not share *Stichodactyla* specialization (*A. akallopisos*–*A. polymnus*, *A. akallopisos*–*A. sebae*, and *A. sandaracinos*–*A. sebae*). In both host categories, the number of SVs shared by specialists falls within the range observed among the control groups, providing no evidence for an excess of shared SVs associated with host specialization.

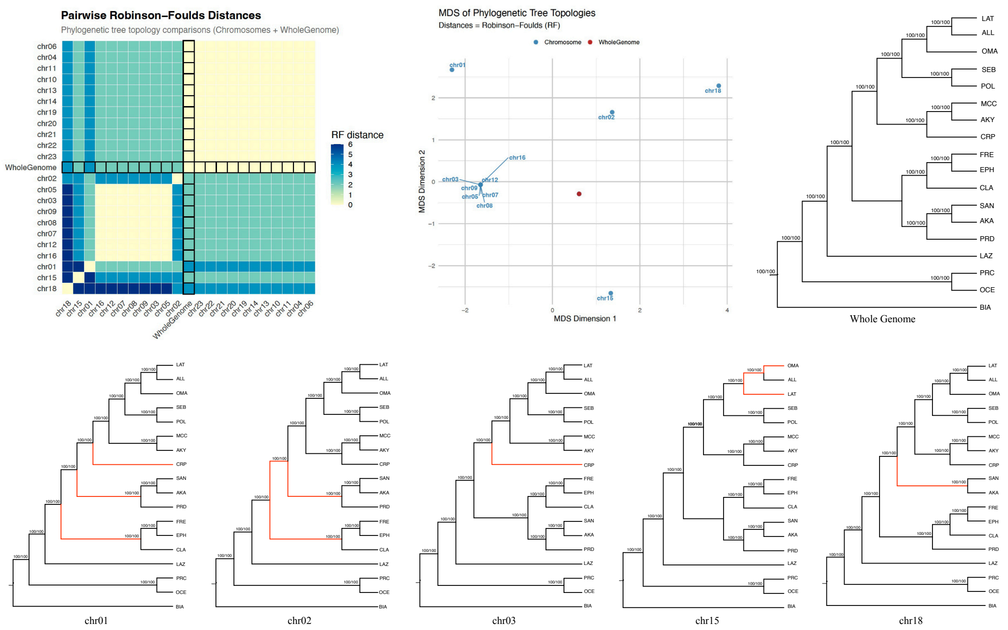


Figure S29: Topological inconsistency across clownfish chromosomes, based on phylogenies inferred from SNPs identified via SyRI alignments to the *A. biaculeatus* reference genome. **Left:** Pairwise Robinson–Foulds (RF) distances among all chromosome-level phylogenies and the whole-genome phylogeny, visualised as a heatmap. Chromosomes with high RF distances relative to the whole-genome tree appear as dark rows and columns. **Centre:** Multidimensional scaling (MDS) of the pairwise RF distance matrix, showing how chromosome-level topologies relate to one another and to the whole-genome topology (red dot). The discordant chromosomes (chr01, chr02, chr03, chr05, chr07, chr08, chr09, chr12, chr15, chr16, and chr18) are labelled. All remaining chromosomes not shown in the MDS plot are positioned with the whole-genome topology. **Right**: Maximum-likelihood tree inferred from the whole-genome concatenated alignment, shown for reference. **Bottom**: Individual maximum-likelihood trees for the five chromosomes forming the most distinct discordance clusters in the MDS (chr01, chr02, chr03, chr15, chr18). Branches deviating from the whole-genome topology are highlighted in red. All remaining chromosomes not shown as individual trees recover the same topology as the whole-genome tree. Node support values indicate SH-aLRT / ultrafast bootstrap.
